# Tumor-specific CD4 T cells cooperate with myeloid cells to remodel the pancreatic tumor microenvironment and enable effective immunotherapy

**DOI:** 10.64898/2026.07.28.741325

**Authors:** Eduardo Cruz-Hinojoza, Zoe C. Schmiechen, Sindri M. Bonner, Adam L. Burrack, Madeline A. Ellefson, Rachana Pandey, Amrit Gaire, Sandhya Appiah, Steven S. Shen, Thamotharampillai Dileepan, Ingunn M. Stromnes

## Abstract

We interrogate antigen-specific CD4 T cells during immunotherapy in pancreatic ductal adenocarcinoma. Vaccination with MHC-II-restricted tumor epitopes impart superior protection compared to an immunodominant MHC-I epitope, prompting development of MHC-II affinity-enhanced tetramers to track tumor-specific CD4 T cells. As tumors progress, tumor-specific CD4 T cells decline, and remaining cells acquire features of regulation. Agonistic anti-CD40 increases Th1 cell clonal expansion and transiently decreases Tregs. Anti-PD-L1 promotes Tfh clonal expansion in draining lymph nodes and tumor while preventing Treg rebound after anti-CD40. Treatment with anti-CD40 promotes intratumoral Stat1^+^ macrophages, tertiary lymphoid structures (TLS), and immune triads. MHC-II on myeloid cells but not B cells is required for immunotherapy-induced TLS formation and antitumor effects. IL-15 complex enhances immunotherapy-induced Th1 effectors without promoting Tregs. Human immunotherapy transcriptomics shows conserved Th1 programming and Treg destabilization as a feature of response. Thus, tumor-specific CD4 T cells are central mediators of effective immunotherapy in solid tumors.

## INTRODUCTION

Pancreatic ductal adenocarcinoma (PDA) is a lethal malignancy, with a five-year survival rate of 8% (1). Despite advances in cancer immunotherapy, PDA remains refractory (2). Resistance is attributed in part to the robust fibroinflammatory and immunosuppressive tumor microenvironment (TME), in which suppressive myeloid populations dominate and impair productive immunity (3).

CD4 T cells are essential components of antitumor immunity (4–6). CD4 T cell activation depends upon recognition of peptide-MHC-II complexes presented by professional antigen-presenting cells (APC). MHC-II-restricted neoantigens are required for optimal antitumor immunity in other tumor settings (7–9). In PDA, MHC-II antigen presentation by Ccr2^+^ myeloid cells is critical for conventional Foxp3-CD4 T cell (Tconv) differentiation into Th1 cells and programming antitumor macrophages (10). Further, CD4 T cell cognate recognition of tumor antigen interferes with intratumoral Foxp3^+^ CD4 regulatory T cells (Treg) accumulation and CD8 T cell exhaustion (10). Thus, reciprocal and cognate interactions between antigen-specific CD4 T cells and myeloid cells can fundamentally reshape the suppressive pancreatic TME at steady state.

Agonistic antibodies to CD40 mimic CD40L, which is upregulated on activated CD4 T cells to and thereby promotes APC activation, and, in cooperation with PD-1/PD-L1 blockade can promote antitumor effects in PDA (11,12). Although, agonistic αCD40, anti-PD-1, and chemotherapy showed a lack of durable benefit in metastatic PDA, increased T_H1_ and T follicular helper (Tfh) CD4 T cells correlated with clinical responses (13,14). It is unknown how these therapies impact tumor-specific CD4 T cells because of the difficulty in identifying the relevant epitopes and tracking such rare CD4 T cells. Unlike CD8 T cells, antigen-specific CD4 T cells can be extremely rare during an immune response, thereby eluding precise analyses. Additionally, there are no known native MHC-II-restricted antigens in preclinical PDA mouse models that could enable faithful analyses of PDA-specific CD4 T cells. Even if an appropriate peptide were identified, MHC-II tetramers are challenging to generate because the peptide requires direct linking to the MHC molecule itself, which requires specialized cloning expertise (15). Further, low affinity CD4 T cells that can contribute to immune responses will often evade detection with traditional peptide:MHC-II tetramers (16).

To address this knowledge gap, we developed an affinity-enhanced MHC-II tetramer to longitudinally track rare endogenous polyclonal tumor-specific CD4 T cells in a PDA orthotopic animal model that faithfully reflects the spectrum of T cell infiltration in human disease (17–19). We show that tumor-specific CD4 T cells adopt distinct fates in the TME yet progressively decline and acquire features of dysfunction while being maintained in tumor draining lymph nodes (dLN) in a Tfh precursor-like state. Intratumoral CD4 T cell clonal expansion was a determinant of antitumor response. Through pMHC-II tetramer staining and image analyses, we demonstrate non-redundant and longitudinal effects of αCD40 and PD-L1 blockade on the tumor-specific CD4 T cells. αCD40 drove the formation of tertiary lymphoid structures (TLS) in tumor periphery, and myeloid:T cell interactions in the tumor core. At the effector phase, therapeutic responses required CD4 T cell licensing of myeloid cells to take up tumor antigen and MHC-II on myeloid cells but not B cells. Despite IL-2Rβ expressed on both Treg cells and Th1 cells, enhancing IL-2Rβ signaling through provision of IL-15 complex during immunotherapy numerically amplified tumor-specific Th1 cells without amplifying their Treg counterparts. Intratumoral reprogramming of CD4 T cell subsets correlates with clinical responses in human clinical trials. Together, our findings uncover a critical CD4 T cell–myeloid cell axis that coordinates pancreatic TME remodeling and promotes effective immunotherapy response, while identifying a translatable therapeutic strategy to amplify this immune circuit.

## RESULTS

### CD4 T cells suppress KPC tumor growth and MHC-II-restricted CB peptide vaccination confers protection

The genetically engineered *Kras*^LSL-G12D/+^;*Trp53*^LSL-R172H/+;^*p48*-Cre (*KPC*) mouse model of invasive PDA expresses the cardinal oncogenic and tumor suppressor driver mutations of human PDA in murine pancreas, mirroring the genetics, histopathological progression, and therapeutic response of human disease (20,21). To investigate CD4 T cell-mediated antitumor mechanisms in a reproducible and rigorous manner, we isolated primary pancreatic tumor epithelial cells from a *KPC* mouse (*KPC*2). After orthotopic implantation into the pancreas of a syngeneic host, *KPC*2 tumors restrict T cell infiltration and are refractory to PD-1/PD-L1 blockade (22). In contrast, *KPC*2 cells transduced with click beetle red luciferase (CB) (*KPC*2a), typically used for bioluminescent imaging, instead results in the accumulation of CB_101-109_:H-2D^b^-specific CD8 T cells in tumors and transient response to PD-1/PD-L1 blockade (22).

To determine if CD4 T cells impact tumor growth in syngeneic *KPC*2 (poorly immunogenic) and *KPC*2a (immunogenic) orthotopic PDA models, CD4 T cells were depleted prior to orthotopic implantation and tumor mass was determined after 14 days. CD4 T cell depletion increased tumor mass in both models (**Figure 1A, B**) indicating globally, CD4 T cells impair tumor growth. As relevant MHC-II antigens are unknown in *KPC* tumors, we hypothesized that the model antigen CBR may provide MHC-II restricted epitopes. We employed I-A^b^ binding prediction algorithms to generate a list of candidate peptides predicted to bind I-A^b^. Top candidate MHC-II-restricted peptides included CB_54-64_ and CB_440-450_ (**Figure 1C; Supplementary Figure 1A**). To determine if these epitopes are processed and presented in tumor-bearing mice, splenocytes isolated from *KPC*2a-bearing mice were restimulated with candidate peptides and IFN-γ-producing T cells were enumerated using ELISpot. CB54 and CB440 peptides elicited IFN-γ-producing T cells (**Figure 1D**). To confirm peptide immunogenicity, we vaccinated B6 mice with peptide in combination with Poly:IC and agonistic αCD40 (TriVax) (22). Seven days later, splenocytes were re-stimulated with corresponding peptides or anchor-substituted peptides in which key MHC-binding residues (P1 or P4) were replaced with either a glycine or glutamine. Wildtype CB_54-64_ and CB_440-450_ peptides, but not altered peptides, elicited IFN-γ production by T cells, demonstrating peptide specificity (**Supplementary Figure 1B, C**).

**Figure 1.**
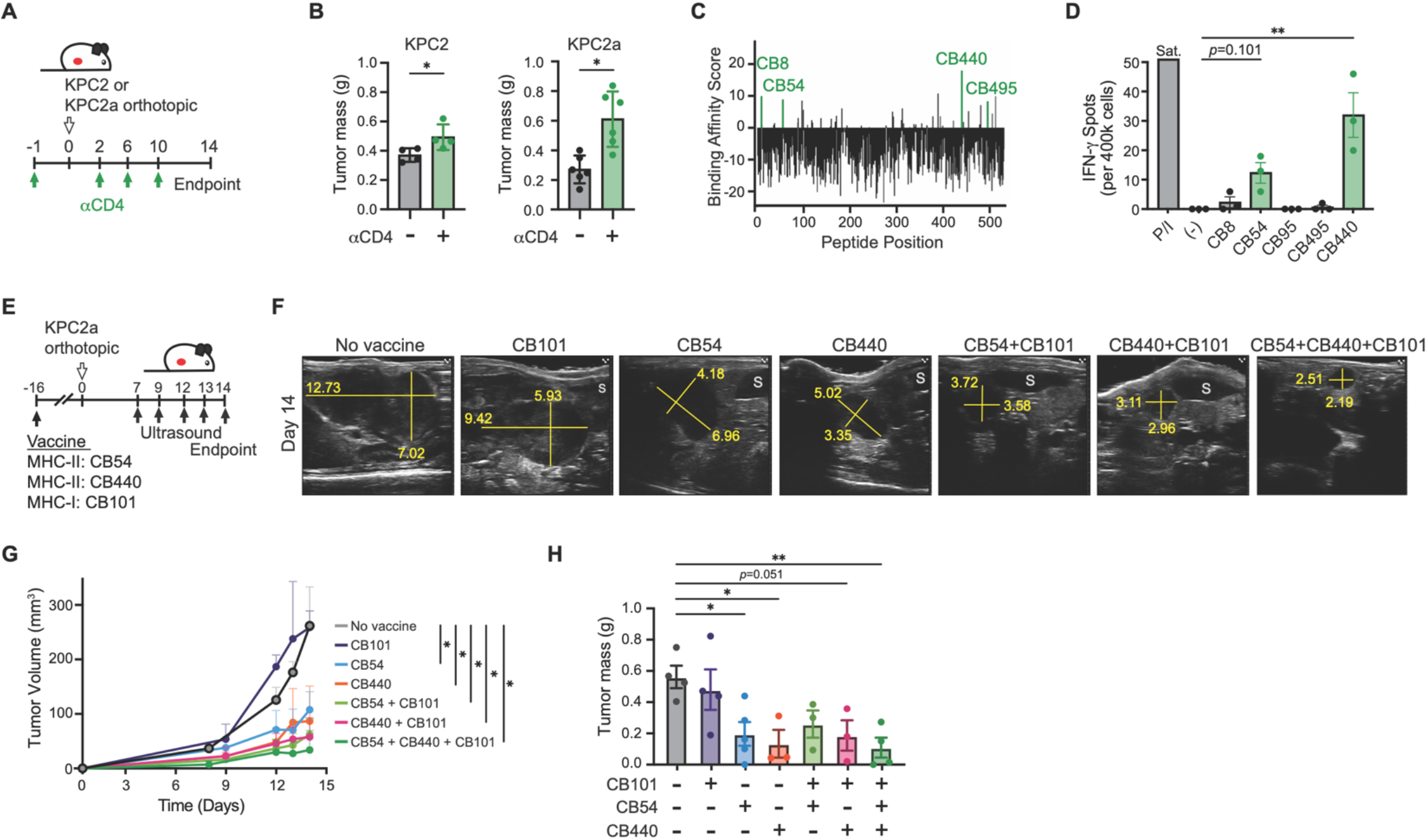
CD4 T cells suppress *KPC* tumor growth and MHC-II-restricted CB peptide vaccination confers protection. (**A**) Schematic of CD4 T cell depletion in orthotopic tumor-bearing mice. (**B**) Tumor weights on day 14 from control and anti-CD4-treated *KPC*2 (left) or *KPC*2a (right)-bearing mice. (**C**) Peptide binding prediction scores to I-A^b^ by peptide position for CB-derived 11mer peptides. Highlighted peptides were selected for experimental validation. (**D**) ELISpot quantification of IFN-γ-producing splenocytes from *KPC*2a-bearing mice restimulated with the indicated peptides. Sat., saturating spot counts. (**E**) Schematic of prophylactic peptide vaccination schedule and *KPC*2a orthotopic tumor implantation. (**F**) Representative ultrasound images on day 14 from vaccinated mice. (**G**) Mean tumor volume over time. (**H**) Tumor weights on day 14. Each dot represents an independent mouse. (B, D, H). Data are mean ± SEM (B, D, G, H). Unpaired student’s T tests (B). One-way ANOVA with Tukey posttest (D, G, H). \**p*<0.05; \*\**p*<0.01

To determine if CB54- or CB440-specific CD4 T cells confer protective antitumor immunity, we prophylactically vaccinated mice with CB54, CB440, or the immunodominant H-2D^b^-restricted epitope CB_101-109_ (22–24). After sixteen days, vaccinated mice were implanted with orthotopic *KPC*2a cells and tumor growth was monitored using high-resolution ultrasound (**Figure 1E**). Vaccination with CB54 or CB440, yet not the immunodominant MHC-I tumor epitope CB101-109, slowed tumor growth (**Figure 1F-G**) and decreased tumor mass at endpoint (**Figure 1H**). Thus, CB54 and CB440 are naturally processed *in vivo*, and polyclonal endogenous CD4 T cells specific to these epitopes confer superior antitumor immunity compared to CD8 T cells specific for a highly immunogenic MHC-I-restricted epitope expressed by the same tumor cells.

### Tracking tumor-specific CD4 T cells using an MHC-II affinity enhanced tetramer

Having established that polyclonal endogenous CD4 T cells specific to CB54 and CB440 epitopes have antitumor effects, we next sought a strategy to track their fate. The tremendous diversity of an endogenous T cell repertoire results in extremely low frequencies of any particular antigen-specific CD4 T cell population, which is estimated to be 0.1-1 in 10^6^ CD4 T cells in the pre-immune repertoire (25). Compared to CD8 T cells, antigen-specific CD4 T cell proliferate less and bind their cognate MHC molecule with lower affinity (16,26–28), making it challenging to track this population even after antigen exposure. As such, we opted to generate an affinity-enhanced MHC-II tetramer, where mutations in the CD4 binding region of the MHC-II beta chain permits detection of both low and high affinity antigen-specific CD4 T cells, thereby increasing sensitivity 2-fold as compared to traditional tetramers (16). While CB440:I-A^b^ failed repeated attempts at tetramer production, we successfully generated CB54:I-A^b^ tetramers. Because antigen-specific CD4 T cells are generally rare, we conjugated the affinity-enhanced CB54:I-A^b^ tetramer to two different fluorophores to minimize detection of false positives. CB54-specific tetramer-binding CD4 T cells were detected in spleens of CB-positive but not CB-negative tumor bearing mice (**Figure 2A**), thereby validating the specificity of this reagent. To determine the naive precursor frequency of CB54-specific CD4 T cells in unmanipulated mice, we employed a tetramer pull-down method to increase the sensitivity of measuring naïve tetramer-specific cells (15). The precursor frequency of CB54-specific CD4 T cells in unmanipulated mice was ∼100 cells, comparable to naïve 2W1S-specific CD4 T cells (25) (**Figure 2B**). CB54-specific CD4 T cells from the preimmunized repertoire were CD44^LOW^, Cxcr5^−^, Cxcr3^−^, and <5% expressed CD25, a surrogate marker for Treg (**Supplementary Figure 2A**). Thus, CB54-specific CD4 T cells in the pre-immune repertoire are primarily naïve and comprise an abundance within an expected physiological range.

**Figure 2.**
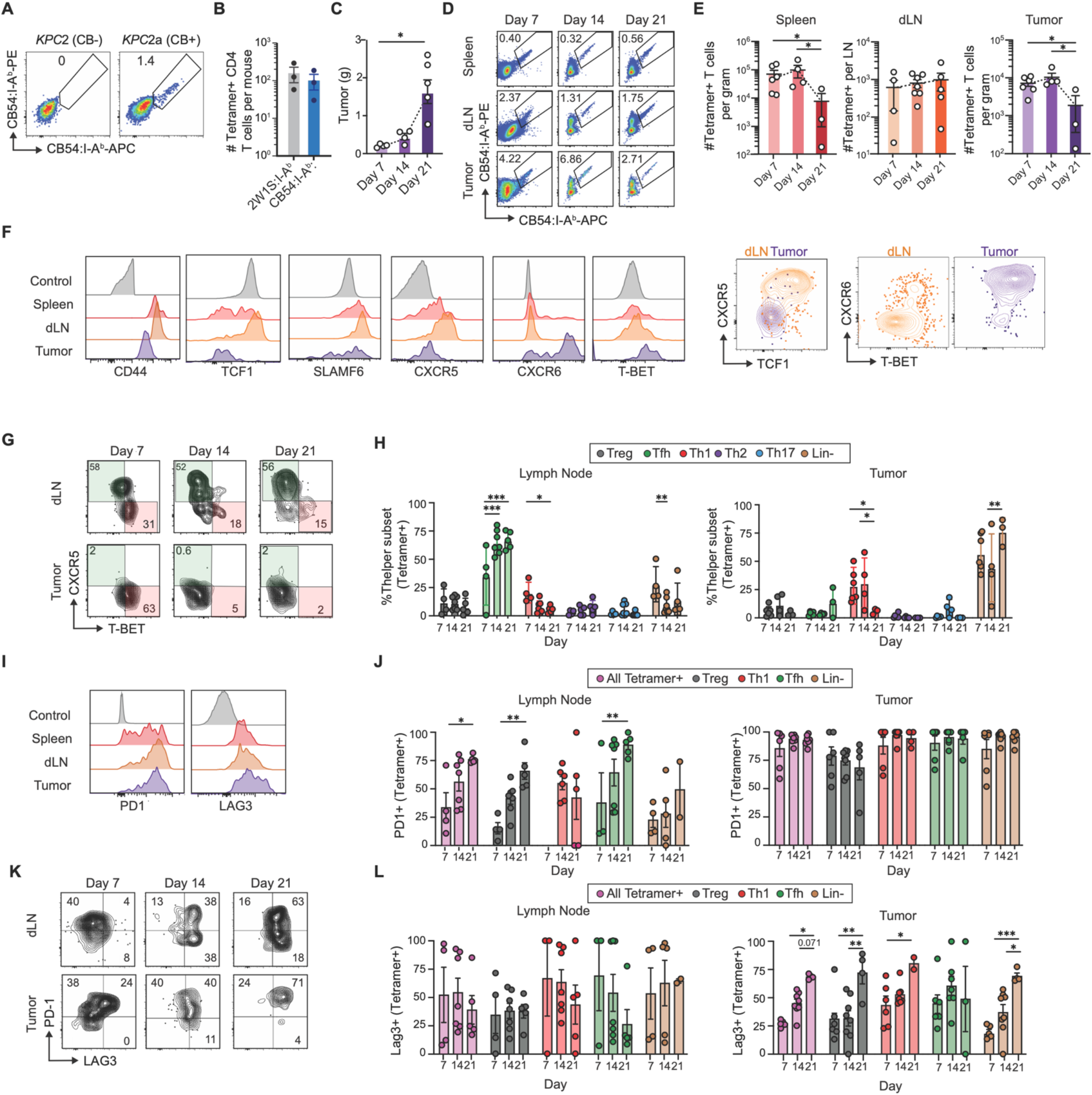
Tracking tumor-specific CD4 T cells using an MHC-II affinity enhanced tetramer. (**A**) Representative flow cytometry plots of CB54:I-A^b^ tetramer staining in *KPC*2 (CB-) and *KPC2a* (CB^+^) tumor-bearing mice. (**B**) Naive precursor frequency of CB54-specific or 2W1S-specific CD4 T cells from pooled secondary lymphoid organs of naïve B6 mice. (**C**) Orthotopic *KPC*2a tumor growth over time. (**D**) Representative flow cytometry plots of CB54:I-A^b^ tetramer^+^ cells gated on viable CD45^+^Dump^-^CD3^+^CD4^+^CD44^+^ cells from spleen, dLNs, and tumor on days 7, 14, and 21. (**E**) Number of CB54:I-A^b^ tetramer binding T cells per gram tumor or spleen, or per dLN. (**F**) Representative staining of CD44, Tcf, Slamf6, Cxcr5, Cxcr6, and T-bet gated on tetramer^+^ CD4 T cells in spleen, dLN, and tumor on day 14. Gray histograms are gated on CD44^−^ CD4 T cells. (**G**) Representative Cxcr5 and T-bet staining gated on tetramer^+^ CD4 T cells in dLNs and tumors over time. (**H**) Th subset composition among tetramer^+^ CD4 T cells in dLNs (left) and tumor (right). (**I**) Representative histograms of PD-1 and Lag-3 among bulk CD44^+^ CD4 T cells (gray) and tetramer^+^ CD4 T cells on day 14. (**J**) Frequency of PD-1^+^ cells among tetramer^+^ subsets in dLNs and tumors. (**K**) Representative plots of PD-1 and Lag-3 gated on tetramer^+^ CD4 T cells in dLN and tumor over time. (**L**) Frequency tetramer^+^ subsets that express Lag-3 in dLN and tumor over time. Data are presented as mean ± SEM, each dot is an individual mouse (B, C, E, H, J, L). One-way ANOVA with Tukey’s posttest (C, E, H, J, L). \**p*<0.05, \*\**p*<0.01, *** *p*<0.001, **** *p*<0.0001.

We sought to characterize CB54:I-A^b^-tetramer binding T cell kinetics during tumor growth by enumerating them on days 7, 14 and 21 post-tumor. Using a two-fluorophore tetramer-staining strategy gated on CD44^high^ CD4 T cells, tetramer^+^ cells comprised <1% in spleen, 1-3% in tumor draining lymph node (dLN), and 2-9% in tumors; a relative frequency that was maintained despite a precipitous increase in tumor growth (**Figure 2C-D**). Although the number of intratumoral CB54-specific CD4 T cells were maintained between day 7 and 14 post-tumor, they decreased 10-fold by day 21, mirroring T cell kinetics in spleen (**Figure 2E**). In contrast, CB54-specific T cell number in the dLN remained steady throughout tumor growth. Although tetramer mean fluorescence intensity (MFI) within the antigen-specific population was similar across tissues (**Supplementary Figure 2B**), we noted marked phenotypic changes occurred within each anatomic location as early as day 14 post-tumor (**Figure 2F**, **Supplementary Figure 2C**). Cxcr5, which defines Tfh in the lymphoid follicle (29,30), was expressed on most tumor-specific CD4 T cells in the dLN, yet was mostly absent in tumors. Tfh in dLN expressed stem-associated markers Tcf1 and Slamf6, paralleling PDA-specific CD8 T cells with stem-like progenitor properties (23). In contrast, Cxcr6, a marker of tissue resident memory T (T_RM_) cells (31), was up-regulated on most tumor-infiltrating tumor-specific T cells but rarely on dLN T cells. Cxcr6 expression tracked with the cardinal Th1 transcription factor T-bet, consistent with a marker of Th1 cells (**Figure 2F**) (6). These data suggest that PDA-specific CD4 T cells are biased toward Tfh cell with stem-like markers in dLN but primarily adopt a more differentiated Th1 state in the pancreatic TME.

To more precisely delineate CB54-specific CD4 T cell differentiation, we employed a sequential gating strategy to distinguish Treg (Foxp3^+^), Tfh (Cxcr5^+^), Th1 (T-bet^+^), Th2 (Gata3+), Th17(Rorγt^+^), and lineage-negative (Lin^−^; Foxp3^−^, Cxcr5^−^, T-bet^−^, Gata3^−^, Rorγt^−^) cells (**Supplementary Figure 2D,** ref. (32)). Expectedly, Cxcr5^+^ tetramer-binding cells in dLN lacked the cardinal Th1 transcription factor T-bet (**Figure 2G**). Tfh in the dLN were also increased in frequency between 7- and 14-days post-tumor (**Figure 2H**). Tfh were rare in tumors (**Figure 2G, H**) and comprised 20-30% of CB54-specific T cells in spleen (**Supplementary Figure 2E**). Approximately 20% of CB54-specific T cells in the dLN were Th1 on day 7 but this population decreased 4-fold by day 21. Additionally, the frequency and number of intratumoral Th1 cells declined between 14 and 21 days, which corresponded with an increase in Lin^−^ cell frequency. In spleen, CB54-specific T cells were mostly Lin^−^ (**Supplementary Figure 2E**). The decline in dLN and intratumoral Th1 cells paired with an increase in the Lin^−^ state suggests that there is a large fraction of tumor-specific CD4 T cells that may be halted in their differentiation trajectory which may be due to competition for antigen and cytokine support.

We next assessed coinhibitory receptors PD-1 and Lag-3 among the dominant subsets (Th1, Tfh, Treg, and Lin^−^), as they were elevated compared to naïve CD4 T cells (**Figure 2I**, gray control). In dLN, PD-1 was progressively upregulated over time, whereas in tumors, PD-1 was induced early and maintained on Treg, Th1, and Lin^−^ cells (**Figure 2I-L**). In tumors and spleen, Lag-3 upregulation increased over time, and most tetramer^+^ CD4 T cells were PD-1^+^Lag-3^+^ by day 21 (**Figure 2I-L, and Supplementary Figures 2F, G**). These changes may be driven to increased antigen, and tumor cell spread to secondary lymphoid organs (6). Further, the co-stimulatory receptor ICOS, known to support CD4 Tconv antitumor responses (33) and maintain Tfh state through suppression of Klf2 (29), progressively declined among Tconv cells yet was maintained on Treg cells (**Supplementary Figures 2H, I**). Together, these data show that tumor-specific CD4 T cell adapt to the local environment and may become impaired during PDA growth.

### αPD-L1 promotes Tfh clonal expansion whereas αCD40 modulates Th subset-specific transcriptional states

Agonistic αCD40 + αPD-L1 (Dual) synergizes to control orthotopically-implanted *KPC*2a tumor growth, yet exerts quite modest antitumor effects in the poorly immunogenic (CBR-negative) *KPC*2 model (11,23). As our prior studies largely focused on tumor-specific CD8 T cells (11), we sought to determine if CD4 T cells were also required for the antitumor effects of Dual immunotherapy. As such, we depleted CD4 T cells 6 days after orthotopic *KPC*2a tumor establishment and one day prior to immunotherapy (**Supplementary Figure 3A**). This delayed CD4 T cell depletion strategy permitted early CD4 T cell priming, licensing of APCs, and their provision of help to CD8 T cells. CD4 T cell depletion a day prior to immunotherapy resulted in larger tumors (**Supplementary Figure 3B**), indicating CD4 T cells are required at the effector phase during immunotherapy.

As αCD40 failed to compensate for a lack of CD4 T cell help, we hypothesized it may instead alter tumor-specific CD4 T cells. As such, we probed scRNA-seq datasets of immune cells in poorly immunogenic (*KPC*2 (6)) or immunogenic (*KPC*2a (23)) orthotopic tumors following αCD40 and/or αPD-L1. Our prior assessment of these datasets focused on CD8 T cells. CD4 T cells were selected based on *Cd3e* and *Cd4* transcripts above the non-zero 1st percentile on day 14 post tumor, which is 7 days post immunotherapy (**Figure 3A**, **Supplementary Figure 3C** (34)). In the poorly immunogenic *KPC*2 model, unsupervised clustering identified naïve (*Cd44*^low^, *Sell*^+^), stem-like (*Tcf7*, *Slamf6*, *Cd44*) (35), Tfh-like (*Tox, Slamf6, Bcl6*) (36), two *Cxcr6*^+^ Th1 clusters, and two Treg cell clusters (**Figure 3B, C, Supplementary Figure 3D**). The most abundant Treg cluster (Treg-1) expressed *Il2ra* and *Ccr8*, receptors enriched on clonally expanded and highly suppressive Treg cells (37). Dual therapy expanded both Th1 clusters and the Tfh cluster, while also decreasing the highly suppressive *Ccr8*^+^ Treg-1 cluster (**Figure 3D**). Both Th1 clusters were enriched for *Cxcr6* and *Tbx21* and coinhibitory receptors (*Pdcd1, Lag3, Ctla4*) (**Figure 3C**). These were distinguished by Th1 effector genes (*Ifng, Gzmk, Nkg7*) supporting distinct functional Th1 states (**Supplementary Figure 3D**).

**Figure 3.**
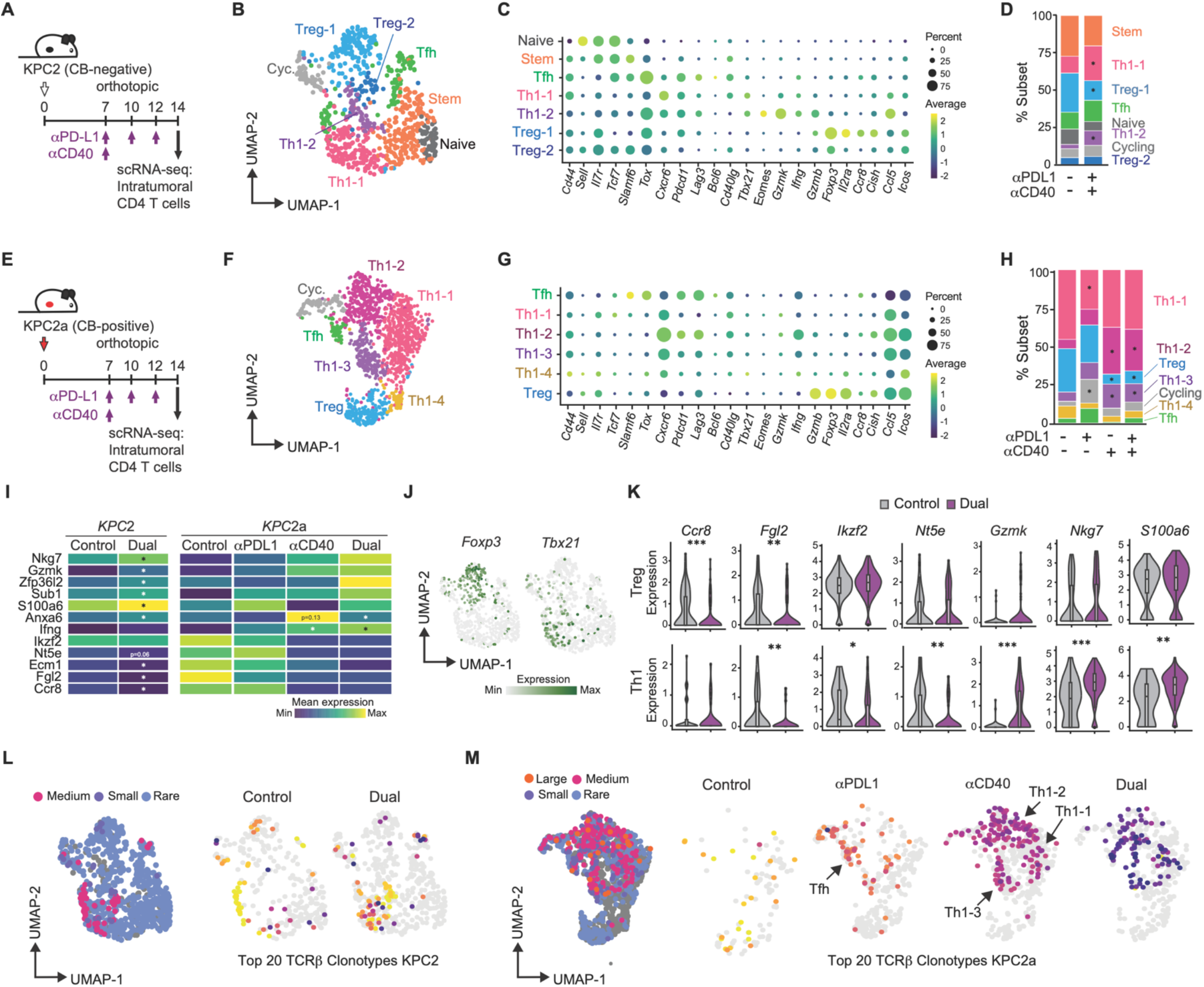
αPD-L1 promotes Tfh clonal expansion and αCD40 modulates Th subset-specific transcriptional states. (**A**) Schematic of immunotherapy treatment and day 14 single-cell capture for orthotopic *KPC*2 (CB^-^)-bearing mice. (**B**) UMAP of intratumoral CD4 T cells in *KPC*2 tumors colored by cluster identity. (**C**) Dot plot of cluster defining genes in (B). (**D**) Stacked bar plot showing cluster frequency per treatment group in *KPC*2 tumors. Asterisks indicate clusters significantly different from control. (**E**) Schematic of immunotherapy treatment and day 14 single-cell capture for T cells in *KPC*2a (CB^+^) tumors. (**F**) UMAP of intratumoral CD4 T cells infiltrating *KPC*2a tumors. (**G**) Dot plot of cluster-driving genes from (F). (**H**) Stacked bar plots showing cluster frequency in *KPC*2a tumors. Asterisks indicate clusters significantly different from control. (**I**) Heatmap of top differentially expressed genes (DEGs) in CD4 T cells in control and treated cohorts from *KPC*2 (left) and *KPC*2a (right) datasets. (**J**) Feature plots of Foxp3 (left) and T-bet (right) in *KPC*2 tumors. (**K**) Violin plots of DEGs in Treg and Th1 subsets. (**L-M**) Feature plots of clonotype size (left) and distribution of top 20 TCRβ clones in CD4 T cells across treatments (right) in *KPC*2 (L) and *KPC2*a (M) (left). Full DEG lists of cluster-driving genes in Supplementary Table 1. Chi-square tests for (D) and (H). Wilcoxon rank-sum tests with Bonferroni correction for (C) and (K). \**p*<0.05; \*\**p*<0.01; \*\*\**p*<0.001.

We next analyzed a four-arm scRNA-seq dataset from *KPC*2a (CBR^+^) tumors from control mice or mice treated with αPD-L1, αCD40, or Dual (23) (**Figure 3E**). Despite performing this experiment at the same timepoint with identical treatment conditions as the poorly immunogenic *KPC*2 model, naïve and stem-like CD4 Tconv cells were undetectable (**Figure 3F, G**). Instead, several *Cxcr6*^+^ Th1 cells dominated in *KPC*2a tumors (**Figure 3F,G**). Paralleling *KPC*2 tumors, a *Il2ra^+^Ccr8*^+^ Treg cell cluster and a *Bcl6*^+^ Tfh-like cluster were identified. Whereas αPD-L1 preferentially expanded cycling cells, αCD40 and Dual expanded multiple Th1 cell clusters that comprised over 80% of the CD4 T cell infiltrate and reduced Treg cells to < 5% of the infiltrate (**Figure 3H, Supplementary Figure 3E**). Consistent with flow cytometry (Figure 2), neither a distinct *Rorc*^+^ Th17 nor a *Gata3*^+^ Th2 cells were undetectable (**Supplementary Figure 3F**).

We next determined immunotherapy-induced differentially expressed genes (DEGs) in CD4 T cells in *KPC*2 and *KPC*2a tumors to garner insight into disparate responses. Although effector genes were upregulated after Dual in both CD4 T cell datasets (*GzmK*, *Nkg7*, *S100a6*), *Ifng* appeared further increased in the *KPC*2a model (**Figure 3I**), likely contributing to disparate responses (11). Immunosuppression-associated genes *Ikzf2* (38), *Fgl2* (39,40), *Ccr8* (*37,41*), and *Nt5e* (CD73) (42) and Tfh-associated gene *Ecm1* (43) were reduced after Dual (**Figure 3I**). αPD-L1 induced proliferating genes in CD4 T cells (**Supplementary Figure 3G**), that were enriched for *Il2* and *Bcl6* (**Supplementary Figure 3H**). These data suggest anti-PD-L1 promotes the proliferation of intratumoral Tfh cells. αCD40 reduced Treg genes (*Il2ra, Foxp3*), increased glycolytic metabolism (*Hif1α, Ldh*), programmed Th1 cells (*Cxcr6, Il12rb2, Il18r1*) and induced myeloid-centric factors (*Csf1, Mif*) (**Supplementary Figure 3G**). We subsetted *Tbx21*^+^ Th1 and *Foxp3*^+^ Treg cells to identify subset-specific transcriptional changes induced by Dual(**Figure 3J**). While *Ccr8* was uniquely decreased in Treg cells, *Fgl2* and *Ecm1* were decreased in both Treg and Th1 cells (**Figure 3K**). *Ikzf2*, typically considered a Treg-specific transcription factor, was strikingly decreased in Th1 cells after Dual therapy (**Figure 3K**). *Gzmk*, *Anxa6*, *S100a6*, and *Nkg7* were upregulated in Th1 cells whereas *Zfp36l2* was induced in Treg cells (**Figure 3K, Supplementary Figure 3I**). By visualizing T cell clonotypes on UMAPs, clonally expanded CD4 T cells were rare in KPC2 tumors regardless of therapy (**Figure 3L, Supplementary Figure 3J, K**). In contrast, αPD-L1 promoted Tfh proliferation whereas αCD40 drove Th1 proliferation (**Figure 3M**). Altogether, αPD-L1 unleashes Tfh cells whereas αCD40 induces subset-specific transcriptional changes to promote Th1 and mitigate Treg cells at the day 14 timepoint.

### αPD-L1 promotes Tfh expansion in dLN, limits intratumoral Treg rebound following αCD40, and increases the Th1:Treg cell ratio

As the above scRNA-seq did not distinguish tumor-specific CD4 T cells, we next interrogated CB54-specific T cells at two timepoints post immunotherapy (**Figure 4A**). All therapies reduced tumor mass, with no significant difference in tumor size between the therapy groups at either timepoint (**Figure 4B**), consistent with prior studies (11,22,23). This is ideal because qualitative differences identified in tumor-specific CD4 T cells will be due to immunotherapeutic interventions rather than tumor size Although αPD-L1 and Dual decreased tetramer^+^ CD4 T cell frequency in dLN, their number increased 10-20-fold (**Figure 4C, D**). In spleen, tetramer^+^ CD4 T cell frequency was d <1% of total CD4 T cells, yet αPD-L1 or Dual increased number on day 21 (**Supplementary Figure 4A**). In tumors, tetramer^+^ T cell frequency was variable, and Dual therapy significantly increased their number 10-fold (**Figure 4E**). While Th17 remained undetectable (**Supplementary Figure 4B**), tetramer^+^ Tfh number increased ten-fold in dLN after αPD-L1 on day 14 (**Figure 4F, G**), and on day 21 when combined with αCD40 (**Figure 4F, G**). Dual therapy increased Th1 number in dLN twenty-fold (**Figure 4G**), resulting in a ten-fold increase in the Th1:Treg cell ratio (**Supplementary Figure 4C**).

**Figure 4.**
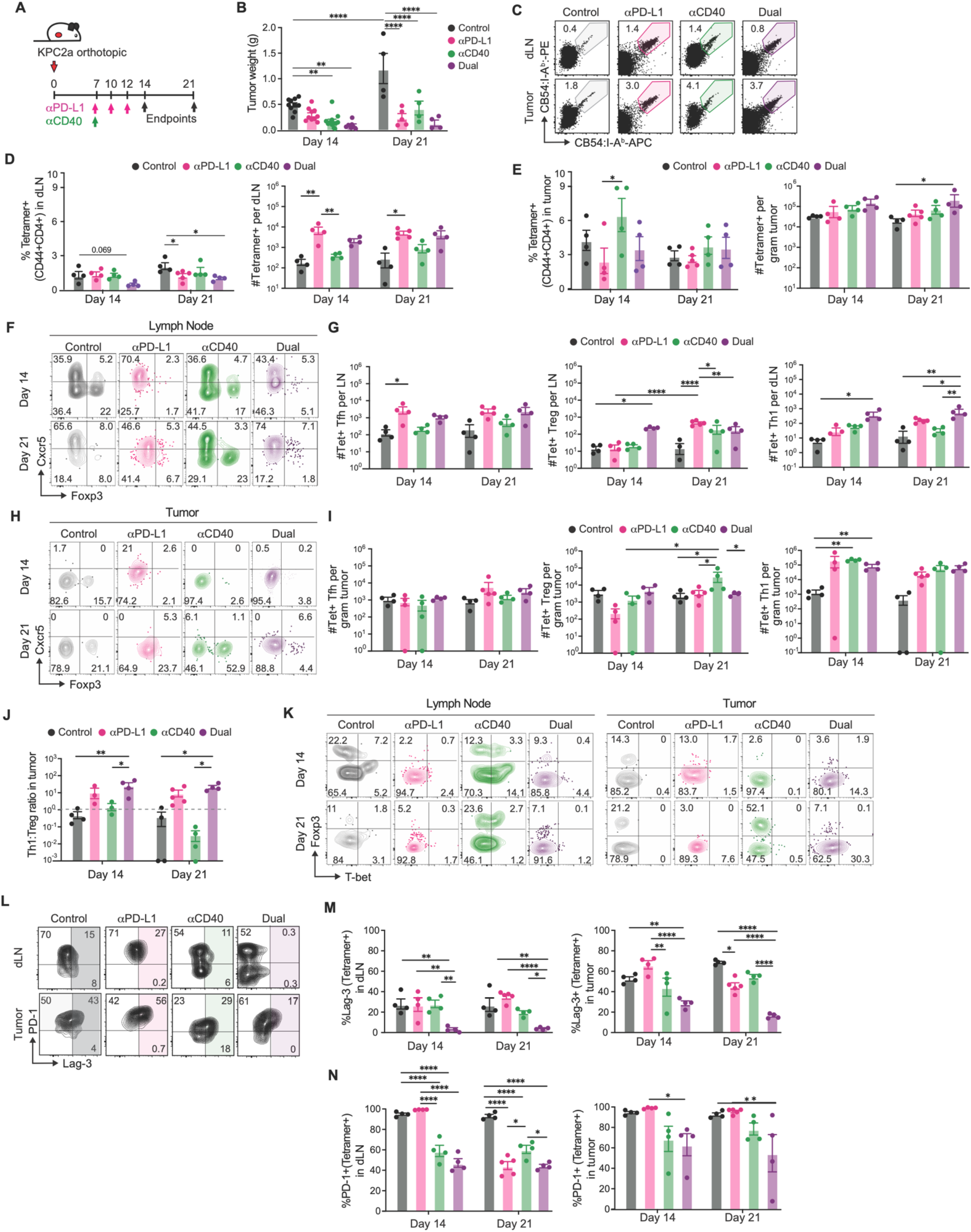
αPD-L1 promotes Tfh expansion in dLN, limits intratumoral Treg rebound following αCD40, and increases the Th1:Treg ratio(**A**) Schematic of treatment schedule. (**B**) Tumor weights on days 14 and 21 across treatment cohorts. (**C**) Representative CB54:I-Ab tetramer staining in dLNs and tumors in each cohort. Cells were gated on live, dump-negative, CD45^+^CD3ε^+^CD4^+^CD44^+^ T cells. (**D** and **E**) Frequency and number of CB54-specific CD4 T cells in the dLN (D) and tumor (E) on days 14 and 21, normalized to tumor weight or LN number. (**F**) Representative Foxp3 and Cxcr5 staining on tetramer^+^ CD4 T cells on days 14 and day 21 across therapies in the dLN, gated as in (C). (**G**) Number of CB54-specific Tfh, Treg, and Th1 cells in the dLN, normalized to LN number. (**H**) Representative plots of Foxp3 and Cxcr5 staining on tetramer+ cells on days 14 and 21, gated as in (C). (**I**) Number of CB54-specific Tfh, Treg, and Th1 cells in tumors on days 14 and 21, normalized to tumor weight. (**J**) Intratumoral Th1 to Treg ratio quantified using numbers calculated in (I). (**K**) Representative plots of Foxp3 and T-bet staining on tetramer^+^ cells on days 14 and 21 in tumor, gated as in (C). (**L**) Representative plots of PD-1 and Lag-3 on tetramer^+^ cells on day 14 in dLN and tumor and gated as in (C). (**M** and **N**) Frequency of Lag-3^+^ (M) and PD-1^+^ (N) cells among CB54 tetramer^+^ cells in dLNs and tumor on days 14 and 21. Data are mean ± SEM, with each dot an independent mouse. One-way ANOVA with Tukey’s postest (B, D, E, G, I, J, M, N). \**p*<0.05, \*\**p*<0.01; \*\*\**p*<0.001; \*\*\*\**p*<0.0001.

In tumors, Tfh frequency (<1% of tetramer-binding CD4 T cells) and number was not impacted by immunotherapies (**Figure 4H-I, Supplementary Figure 4B**). All immunotherapies decreased Treg cell frequency at day 14, yet αCD40 induced a tenfold rebound in tetramer^+^ Treg cells in tumors by day 21 (**Figure 4H-I, Supplementary Figure 4B**), thereby decreasing the intratumoral Th1:Treg ratio (**Figure 4J**). αPD-L1 mitigated the Treg rebound after αCD40 (**Figure 4I, Supplementary Figure 4C**). In inflamed, Th1-polarizing conditions, Treg cells upregulate T-bet that promotes Treg cell proliferation (44). In dLNs, 20% of tetramer^+^ Tregs co-expressed T-bet in control and αCD40-treated animals yet T-bet was absent in tetramer^+^ Treg cells following αPD-L1 (**Figure 4K, Supplementary Figure 4C**). Thus, T-bet in Treg cells following αCD40 may contribute to the T_REG_ cell rebound in tumors (44), or reflect Treg cell conversion to Th1 (45). Although αCD40 alone was sufficient to reduce PD-1, the fraction of tetramer-binding cells expressing Lag-3 was distinctively reduced by Dual in dLN and tumor (**Figure 4L-N**).

### αCD40 therapy induces intratumoral tertiary lymphoid structures formation and immune cell triads

We performed multiplex immunofluorescence to determine if immunotherapies impacted spatial organization of immune cells (**Figure 5A**). Although all therapies increased CD4 T cell infiltration into the tumor core, αCD40 and Dual effects was most dramatic (**Figure 5A-C, Supplementary Figure 5A, B**). Dual therapy reduced tumor cytokeratin (CK) staining (**Supplementary Figure 5A-C**), consistent with more durable antitumor effects (11). αCD40 and Dual induced B cell rich immune aggregates resembling tertiary lymphoid structures (TLS) (**Figure 5B**), organized lymphoid formations found in non-lymphoid tissues in response to chronic inflammation, infection, and cancer (46–48) including human PDA (17,49). Immunotherapy-induced TLS contained a dense array of CD4 T cells, B cells, CD8 T cells, macrophages, and dendritic cells (DCs), and exhibited abundant MHC-II staining (**Figure 5B**). TLS were primarily on the interface between healthy pancreas and primary tumor (peritumoral), suggestive of a barrier that restrains the tumor growth (**Supplementary Figure 5A**). The therapies did not increase B cells, CD8 T cells, or myeloid cell infiltration into the tumor core (**Figure 5C-F**). Both the number of TLS and TLS size at the peritumor border were significantly increased after αCD40 or Dual therapy (**Figure 5G, H**).

**Figure 5.**
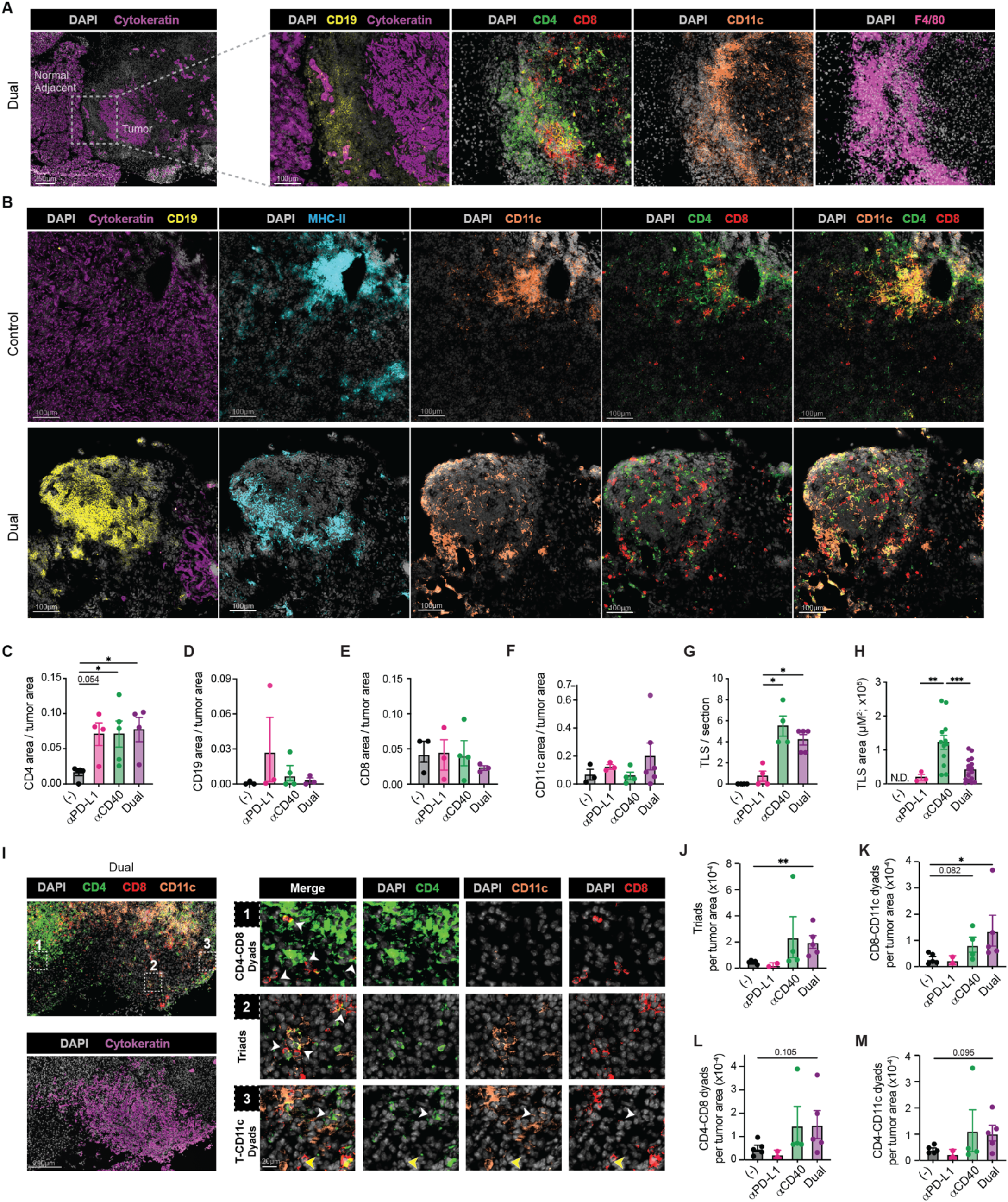
αCD40 therapy induces intratumoral tertiary lymphoid structures and immune cell triads. (**A**) Representative multiplex immunofluorescence images of orthotopic *KPC*2a tumors from a Dual therapy treated mouse showing normal adjacent pancreas, peritumoral region, and intratumoral regions identified by cytokeratin (CK) tumor cell staining (left). Magnified inset shows shows tertiary lymphoid structures (TLS) in tumor periphery. (**B**) Representative multiplex immunofluorescence orthotopic *KPC*2a tumor sections from control or Dual-therapy treated mice and stained for cytokeratin (CK), MHC-II, CD11c, CD4, and CD8α. (**C-F**) Quantification of CD4 (C), CD19 (D), CD8 (E), or CD11c (F) immune cells in tumor core. (**G** and **H**) TLS number per tumor section (G) and TLS area (H). N.D., none detected. (**I**) Representative multiplex immunofluorescence images illustrating immune proximity quantification in orthotopic *KPC*2a tumor from a Dual therapy treated mouse. Left: overview of CD4, CD8, CD11c, and DAPI staining with numbered ROIs (top) and corresponding CK staining (bottom). Right: magnified ROIs showing representative examples of CD4-CD8-CD11c triads and CD4-CD8, CD8-CD11c, and CD4-CD11c dyads indicated by arrows. (**J-M**) Quantification of CD4-CD8-CD11c triads (J), CD8-CD11c dyads (K), CD4-CD8 dyads (L), and CD4-CD11c dyads (M) measured and normalized to tumor area (tumor core + peritumoral region) across treatment groups. Data are mean ± SEM and each dot is an independent mouse (C, D, E, F, G, H, J, K, L, M). One-way ANOVA with Tukey’s multiple comparison correction (C, D, E, F, H, J, K, L, M). For G, data determined to be log-normally distributed by the Shapiro–Wilk test were analyzed by log-normal one-way ANOVA, \**p*<0.05; \*\**p*<0.01; \*\*\**p*<0.001.

The spatial organization of three-cell hubs composed of a CD8 T cell, CD4 T cell, and CD11c myeloid cell is associated with immunotherapy response in melanoma and colon cancer models (9). Immune triads have been associated with clinical immunotherapy response in humans (9) and are associated with T cell clonal expansion prolonged survival in human PDA (50). Within the tumor core and periphery, we identified closely associated (<20 μm) clusters of CD4 T cells, CD8 T cells, and CD11c APCs (triads, **Figure 5I**). Whereas Dual therapy significantly increased the frequency of intratumoral triads, and dyads composed of a CD8 T cell and a CD11c^+^ APCs, dyads composed of CD4-CD8 or CD4-CD11c cells were unchanged (**Figure 5J-M**).

Both TLS and triads are likely spatial areas where CD4 T cells recognize cognate tumor antigen (48,51). To determine if CD4 T cells impacted tumor antigen uptake by myeloid cells, we depleted CD4 T cells and measured the tumor marker GFP in myeloid cells using the gating strategy in (**Supplementary Figure 5D**). CD4 T cell depletion reduced tumor GFP staining in macrophages and dendritic cells (**Supplementary Figure 5E**). Thus, APC tumor antigen uptake may be dependent on CD4 T cells.

As TLS were enriched for B cells, we asked if B cells were necessary for antitumor effects of Dual. However, B depletion did not abrogate antitumor effects of Dual therapy at day 14 (**Supplementary Figure 6A-C**). Similarly, genetic loss of MHC-II only on B cells also did not abrogate the antitumor effect of Dual therapy (**Supplementary Figure 6D-F**). Thus, while B cells are abundant in immunotherapy-induced TLS, they were dispensable for antitumor effects.

### Immunotherapy transforms intratumoral myeloid cells and myeloid MHC-II is required for TLS formation and immunotherapy response

As B cells were dispensable for antitumor effects, we considered myeloid cells may instead be critical. Intratumoral mononuclear phagocytes from immunotherapy-treated mice with KPC2a tumors were subsetted based on *Itgam*^+^*H2-Ab1*^+^ and negative for *Trac,*, *Cd19, and Csf3r*^-^ (**Figure 6A-D, Supplementary Figure 6G, H**). The most abundant cluster was enriched for *Stat1,Cxcl9*, and *Cxcl16* and increased proportionally after αPD-L1, αCD40 and Dual (**Figure 6D**). *Nos2*^+^*Arg1*^+^ cells (C1) were modestly decreased by αPD-L1 yet increased by αCD40. Strikingly, *Ccr2*^+^ monocytes (C2) were decreased by all therapies suggesting that immunotherapy may drive differentiation, consistent with a role for CD4 T cells in modulating monocyte fate in PDA at steady state (10). Two *Apoe*^+^ macrophage clusters (C3, C4) were enriched for *Trem2*, *Maf*, *Ccl8* and distinguished by *Folr2* (**Figure 6D, Supplementary Figure 6H**). Dual therapy distinctively increased the *Folr2*-enriched subset (**Figure 6D**). Although αCD40 can activate dendritic cells (DC), we found *Ccr7*^+^ *Cd274*^+^ *Cd200*^+^ mature immunoregulatory DCs (C5; mregDC) (52), and *Xcr1*^+^ *Zbtb46*^+^ type I DCs (C7; cDC1s) were rare in PDA and reduced in frequency after αCD40 or Dual (**Figure 6D, Supplementary Figure 6H, Supplementary Table 2**).

**Figure 6.**
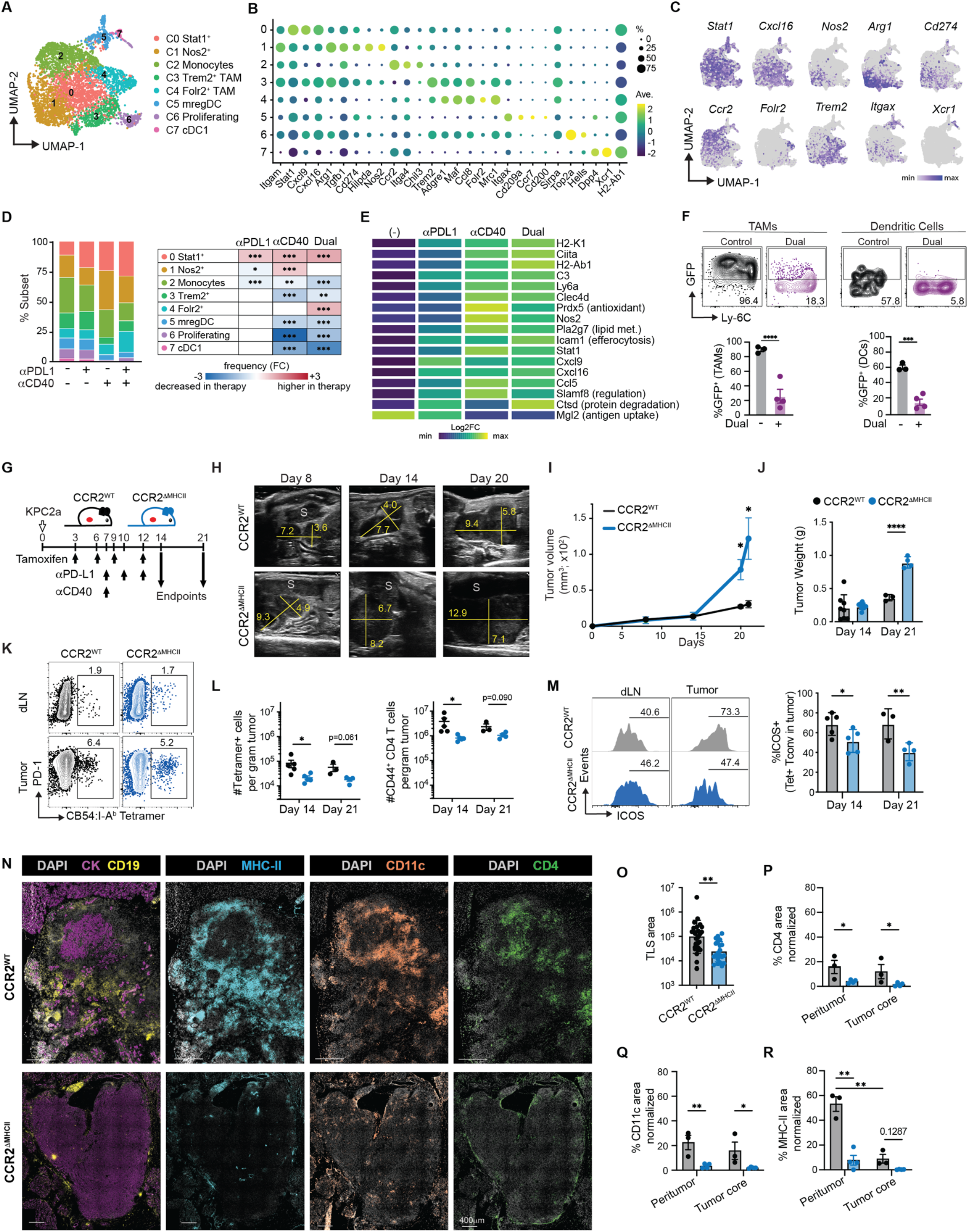
Immunotherapy transforms intratumoral myeloid cells and myeloid MHC-II is required for TLS formation and immunotherapy response. (**A**) UMAP of intratumoral myeloid cells from *KPC*2a tumor-bearing mice colored by cluster identity. (**B**) Dot plot of select cluster-defining marker genes from (A). (**C**) Feature plots of key marker genes. Projected UMAP space from (A). (**D**) Stacked bar plot of myeloid cluster frequencies across treatment groups (left) and heatmap of fold change difference in frequency of each myeloid cluster compared to control group (right). (**E**) Heatmap of selected DEGs between each therapy and control across myeloid clusters. DEGs were identified by Seurat FindMarkers. (**F**) Representative plots of tumor antigen uptake (GFP+) by tumor-infiltrating macrophages or DCs (gating as in Figure S6J) from control or Dual-treated mice (top) and percent myeloid cell positive for GFP (**G**). Schematic of tamoxifen administration and treatment schedule followed and assessed on days 14 and 21 in CCR2^WT^ or CCR2^ΔMHCII^ mice with *KPC*2a orthotopic tumors. (**H**) Representative ultrasound images over time from Dual-treated CCR2^WT^ and CCR2^ΔMHCII^ mice. (**I**) Tumor volume in CCR2^WT^ and CCR2^ΔMHCII^ mice from (H). (**J**) Tumor weights on days 14 and 21 in CCR2^WT^ and CCR2^ΔMHCII^ mice. (**K**) Representative CB54:I-A^b^ tetramer staining in tumors from Dual therapy-treated CCR2^WT^ and CCR2^ΔMHCII^ mice and gated on CD4 T cells. (**L**) Number of tumor-infiltrating CB54-specific tetramer^+^ CD4 T cells (left) and bulk CD44^+^ CD4 T cells (left) in CCR2^WT^ and CCR2^ΔMHCII^ mice on days 14 and 21. (**M**) Representative ICOS staining gated on tetramer^+^ CD4 T cells in the dLN and tumor (left) and frequency of tetramer^+^ CD4 T cells that express ICOS on days 14 and 21 (right). (**N**) Representative multiplex immunofluorescence images of tumor sections from Dual-treated CCR2^WT^ and CCR2^ΔMHCII^ mice. Scale bar = 400 μm. (**O**) TLS area in CCR2^WT^ and CCR2^ΔMHCII^ mice. (**P**-**R**) MHC-II (O), CD4 (P), and CD11c (Q) area normalized to either peritumoral or tumor core regions. Data are presented as mean ± SEM, each dot a mouse. Full DEG results for (D) and (E) cluster differentiation and cross-treatment comparison in Supplemental Table 2. Chi-square test (D), Wilcoxon rank-sum tests with Bonferroni correction (Seurat FindMarkers) (E), one-way ANOVA with Tukey’s multiple comparison correction for (I, J, L, M, P-R). Unpaired Student’s t test (O). \**p*<0.05; ** *p*<0.01; \*\*\**p*<0.001.

We next assessed DEGs and conducted gene set enrichment analysis to identify how immunotherapy impacts myeloid cell functionality. αCD40 and Dual markedly increased antigen presentation genes (**Figure 6E, Supplementary Figure 6I**). Dual therapy also increased myeloid expression of chemokines including *Cxcl9* and *Cxcl16* which bind Cxcr3 and Cxcr6 expressed on antigen-experienced T cells (Figure 3). Dual increased *Nos2* (iNOS) which can induce tumor cell death (53,54), *Icam1* which promotes efferocytosis (55), and *Ctsd* (cathepsin D) involved in protein degradation. αCD40 decreased *Mgl2* that is involved in receptor-mediated endocytosis and tumor antigen uptake (56). We quantified tumor antigen internalization by assessing the tumor marker GFP in myeloid cells following Dual therapy. Dual therapy reduced tumor-derived GFP in TAMs and DCs (**Figure 6F**). Thus, αCD40 either reduces tumor antigen uptake or increases antigen processing by myeloid cells.

To infer how CD4 T cell signals shape macrophage response following immunotherapy, we employed NicheNet (57) with CD4 T cells as the sender cells and myeloid cells as receiver cells. We selected genes that were differentially expressed on myeloid cells following Dual therapy. CD4 T cell derived *Tnf* and *Ifng* were the highest predicted regulators of macrophage transcriptional changes following immunotherapy, with TGF-β (*Tgfb1*), RANKL (*Tnfsf11*), and *Csf1* also ranking highly (**Supplementary Figure 6J**). CD4 T cell derived *Tnf* and *Ifng* were predicted to polarize myeloid cells towards an antitumor phenotype through upregulation of antigen presentation (*Ciita, H2.D1, H2.K1*), co-stimulation (*Cd40*), chemokines (*Ccl5*, *Cxcl10, Cxcl2*), inflammatory molecules (*Tnfa*, *Il1a*, *Nos2),* and genes critical for myeloid cell retention (*Icam1*, *Vcam1*, *Cxcr4*). T cell-derived *Csf1* induced *Tnf* and *Il1a* in myeloid cells and increased lipid metabolism genes (*Nr1h3*, *Pltp*). RANKL was predicted to promote myeloid survival (*Birc3*, *Tnfaip3*). Dual-therapy-induced *Tgfb1* in CD4 T cells was linked to immunosuppression genes (*Serpine1*, *Trem2*, *Hmox1*). Together, these data suggest CD4 T cells deliver complex and potentially competing signals to myeloid cells.

To investigate if immunotherapy depends on CD4 T cell recognition of tumor antigen presented by myeloid cells, we employe a genetic approach. CCR2-Cre^ER^-GFP×MHC-II^fl/fl^ (CCR2^ΔMHCII^) mice with orthotopic *KPC*2a tumors were treated with tamoxifen to delete MHC-II only on Ccr2+ myeloid cells. Tamoxifen was also administered to control MHC-II^fl/fl^ (CCR2^WT^) littermates to control for tamoxifen-induced immunological effects. To validate this model, we developed a gating strategy to assess immune subsets, including DCs, without using MHC-II (**Supplementary Figure 7A**). Tamoxifen induced loss of MHC-II and expression of Cre recombinase (GFP+) specifically on myeloid cells in CCR2^ΔMHCII^ but not CCR2^WT^ mice, consistent with myeloid-restricted Cre inducing MHC-II deletion (**Supplementary Figure 7B**). To leave CD4 T cell priming intact, tamoxifen was given on day 3 posttumor, followed by every 3 days after that to maintain loss and both cohorts were treated with Dual therapy (**Figure 6G**). High-resolution ultrasound showed similar tumor volume at day 14, but by day 21, CCR2^ΔMHCII^ mouse tumors rapidly progressed (**Figure 6H, I**). Tumor mass was increased in CCR2^ΔMHCII^ mice at day 21 (**Figure 6J**). Tetramer^+^ CD4 T cells were readily detected in dLN and tumors in both cohorts (**Figure 6K**). However, MHC-II deletion on myeloid cells reduced tetramer^+^ CD4 T cell number and total antigen-experienced CD4 T cell number in tumors (**Figure 6L**). Additionally, ICOS upregulation on intratumoral tumor-specific CD4 T cells was dependent on MHC-II on myeloid cells (**Figure 6M, Supplementary Figure 7C**). Together, these data indicate antigen presentation by myeloid cells in the TME is critical for CD4 T cell-mediated antitumor effects during immunotherapy. *KPC*2 and *KPC*2a tumor cells readily upregulated MHC-I after IFN-γ, MHC-II staining was not detected (**Supplementary Figure 7D**), excluding a requirement for direct tumor cell presentation to CD4 T cells for immunotherapy effects.

We next assessed the spatial consequences of MHC-II ablation on CCR2^+^ myeloid cells. TLS-like structures were reduced in number and size in CCR2^ΔMHCII^ tumors as compared to control tumors (**Figure 6N, O,** and **Supplementary Figure 7E**). As ICOS promotes TLS formation (58), antigen-recognition by myeloid cells may be critical for the initiation of TLS. We measured T cells in peritumoral regions (<100 μm from tumor border (59)) and tumor core (**Supplementary Figure 7F**). CD4 T cell density was reduced in both the tumor core and peritumoral region when MHC-II was deleted on myeloid cells (**Figure 6P**), while CD8 T cell changes were minimal (**Supplementary Figure 7G,H**). CD11c^+^ cell density and MHC-II staining was markedly reduced in both the peritumor and tumor core in the absence of MHC-II on myeloid cells (**Figure 6N, Q, R**). MHC-II was also not detected on tumor cells *in situ*, further supporting that the antitumor mechanism is indirect (**Figure 6N**). Altogether, these data establish that myeloid cell antigen presentation to CD4 T cells is critical for their penetration into the tumor core, the generation of TLS-like niches, and for antitumor effects of immunotherapy.

### IL-15C enhances immunotherapy-driven tumor-specific CD4 T cell effector differentiation and function

To identify how immunotherapy impacts CD4 T cells in human solid tumors, we interrogated a scRNA-seq dataset patient on head and neck squamous cell carcinoma (HNSCC) and enrolled in a neoadjuvant clinical trial with anti-PD-1 (nivolumab; Nivo), Nivo + anti-CTLA-4 (ipilimumab; Ipi), and Nivo + anti-LAG-3 (relatlimab; Rel) (60). Unsupervised clustering of CD4 T cells revealed two Treg cell clusters (C0, C3), one of which was more activated (*CTLA4*^+^, *IL2RA*^+^;C3), as well as naive (*TCF7*^+^*CD44*^LOW^;C1), Th1 (*GZMK*^+^*CXCR6*^+^ *LAG3*^+^;C2), memory (*CD44*^high^*FOXO1*^+^, C4, (61)), and proliferating cell clusters (**Figure 7A,B**). A fraction of Th1 cells co-expressed mRNA and protein for CD8 and were enriched for *NKG7*, and tissue residency genes (*ITGAE, CXCR6, CD39*) (**Supplementary Figure 8A**). All three therapeutic regimens increased Th1 frequency and a subset of shared genes (**Figure 7C, Supplementary Figure 8B**), yet only Nivo + Ipi decreased *Foxp3* and activated Treg cells (**Figure 7C, D**). Nivo or Nivo + Rel increased Th1 genes *CXCR6*, *IL2RB, TOX2, and IFNG* as well as *IL2RB* (**Figure 7D**). Broadly, a stem-like progenitor gene signature (62) and the hallmark stem-associated transcription factor *TCF7* was reduced post-treatment (**Figure 7D, Supplementary Figure 8C**). As TLS share features of germinal centers (GC), we employed a GC formation score and found it was elevated in CD4 T cells in response to all treatments (**Supplementary Figure 8C**), suggesting orchestration of TLS.

**Figure 7.**
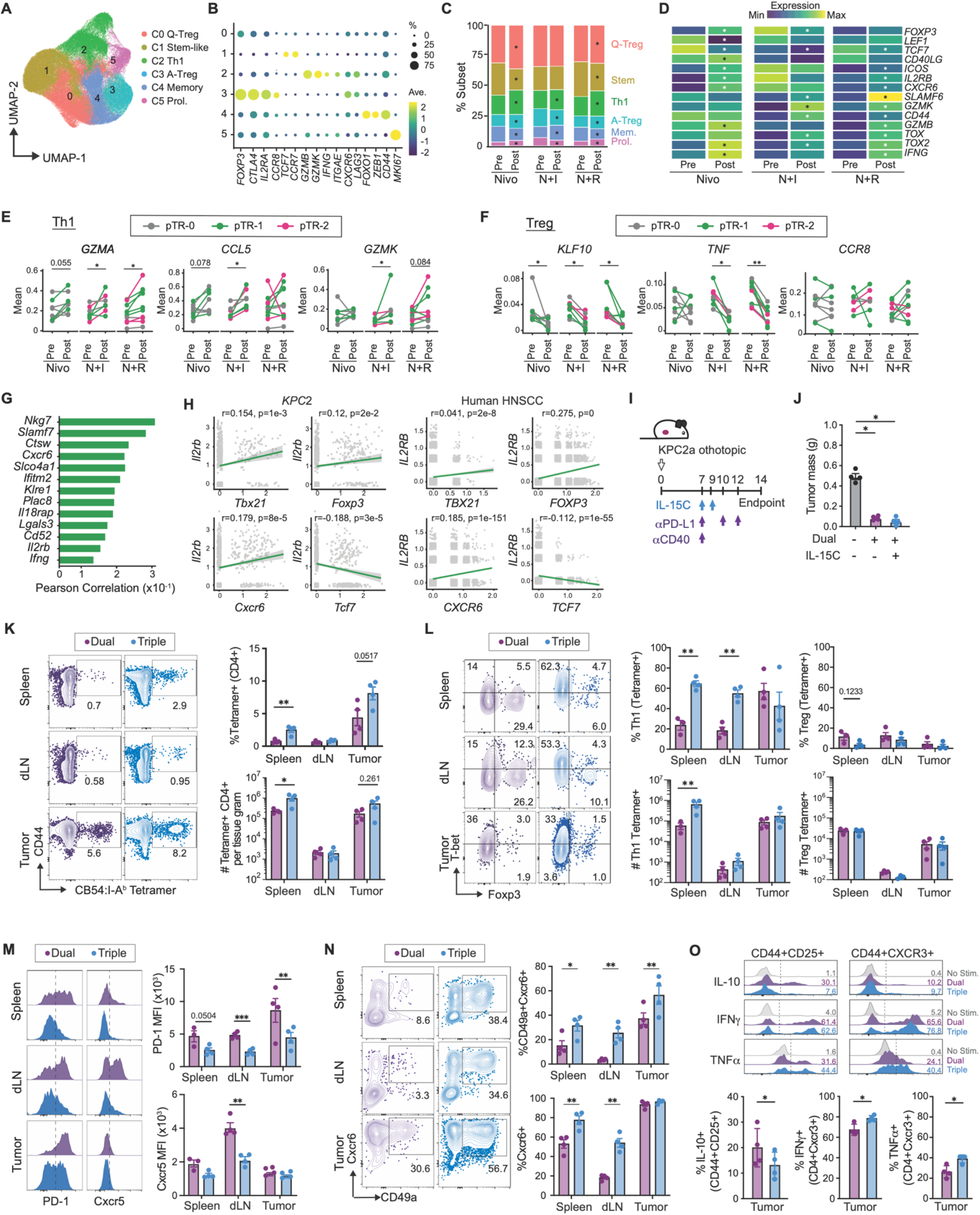
IL-15C enhances immunotherapy-driven tumor-specific CD4 T cell effector differentiation and function. (**A**) UMAP of CD4 tumor-infiltrating T cells from HNSCC patients colored by cluster identity. (**B**) Dot plot of cluster-defining marker genes from (A). (**C**) Stacked bar plots showing cluster frequencies pre- and post-treatment within each treatment arm (nivolumab [Nivo], nivolumab + ipilimumab [N+I], and nivolumab + relatlimab [N+R]). Asterisks indicate cluster is statistically different post-therapy administration. (**D**) Heatmap of select DEGs between pre- and post-treatment CD4 T cells split by treatment arms. (**E** and **F**) Paired pre- and post-treatment expression of select effector genes in Th1 (E) and Treg (F) subsets per patient, split by treatment arm, colored by histopathological response. (**G**) Pearson correlation coefficients of top genes co-expressed with *Tbx21* in CD4 intratumoral T cells from the *KPC*2 dataset. (**H**) Scatter plots of *IL2RB* and select genes in CD4 T cells from *KPC*2 (left) and in human HNSCC (right). Pearson correlation coefficients (r) and BH-adjusted p values are shown with trendline in green. (**I**) Schematic of Dual and Triple (Dual + IL-15C) treatment schedule in orthotopic *KPC*2a-bearing mice. (**J**) Tumor weights at endpoint (day 14). (**K**) Representative plots of CB54:I-A^b^ tetramer staining gated on CD4 T cells (left) and quantification of tetramer^+^ frequency among CD4 T cells (top right) and tetramer^+^ cell number per tissue gram (bottom right). (**L**) Representative plots gated on tetramer+ cells showing T-bet and Foxp3 staining (left), Th1 frequency and number (middle), and Treg frequency and number (right). (**M**) Representative histograms of PD-1 and Cxcr5 gated on tetramer^+^ CD4 T cells across tissues (left) and MFI quantification (right). (**N**) Representative CD49a and Cxcr6 staining in tetramer^+^ CD4 T cells across tissues (left) and percent of CD49a^+^Cxcr6^+^ (top right) or Cxcr6^+^ (bottom right) among tetramer^+^ CD4 T cells. (**O**) Representative histograms of IL-10, IFN-γ, and TNF in Treg (CD44^+^CD25^+^) and Th1 (CD44^+^CXCR3^+^) tumor-infiltrating cells from Dual or Triple therapy-treated mice following PMA/ionomycin restimulation (top) and quantification (bottom). Gray shows unstimulated (No Stim.) cells. Full DEG results for cluster differentiation (B) and cross-treatment comparison (C) are in Supplementary Table 3. Data are mean ± SEM, each dot is an independent mouse (J, K, L, M, N, O).. Wilcoxon rank-sum test with Bonferroni correction (Seurat FindMarkers) (D). One-way ANOVA with Tukey’s posttest (J-N). Unpaired Student’s T-test (O). \**p*<0.05; \*\**p*<0.01, \*\*\**p*<0.001, and \*\*\*\**p*<0.0001.

To directly test whether pre-existing CD4 T cell transcriptional programs associate with therapeutic benefit, we stratified patients by pathological tumor response (pTR-2 responders versus pTR-0/pTR-1 non-responders) in the Nivo+Ipi and Nivo+Rel cohorts (**Supplementary Figure 8D**). There were too few responders in Nivo alone group to pursue such an analysis (60). Responders harbored significantly lower *FOXP3* and *CTLA4* prior to therapy in both Nivo+Ipi and Nivo+Rel cohorts (**Supplementary Figure 8E**), suggesting that Treg cell enrichment in the TME may impair immunotherapy. Additionally, responders in the Nivo+Rel cohort expressed higher levels of the cytotoxic effector molecules *GZMM* and *GNLY* before treatment, suggesting that a pre-existing cytotoxic CD4 Th1 program may contribute to pathological response.

To determine subset specific transcriptional changes, we selected Th1 cells using a Th1 gene module and Tregs using *FOXP3* (**Supplementary Figure 8F, G**). Th1 cells upregulated *GZMA*, *CCL5*, and *GZMK* post-treatment across most arms (**Figure 7E**). Treatment reduced Treg cells expression of *KLF10*, a TGF-β responsive transcription factor known to promote *FOXP3* (63), and *TNF,* which stabilizes Treg cell fate through binding TNFR2 (64) (**Figure 7F**). *CCR8* in Treg cells remained unchanged (**Figure 7F**), suggesting the combination immunotherapies may partially interfere with Treg cells but are insufficient to eliminate the most suppressive state. Thus, immune checkpoint therapies promote Th1 while destabilizing Treg cells in human solid tumors, paralleling our mouse studies.

To identify a strategy to promote tumor-specific Th1 cells during Dual therapy, we analyzed our *KPC*2 tumor scRNA-seq CD4 T cell dataset and identified the top genes associated with the cardinal Th1 transcription factor *Tbx21* (**Figure 7A**). *Il2rb* was among the top 10 genes positively correlated with *Tbx21* (**Figure 7G**). In both human and mouse tumor-infiltrating CD4 T cells, *Il2rb* was also positively correlated with Th1 marker *Cxcr6*, Treg marker *Foxp3*, and negatively correlated with naïve/stem/memory marker *Tcf7* (**Fig 7H**). IL-2Rβ is a shared receptor subunit for mediating IL-2 and IL-15 cytokine signaling. We previously showed that provision of IL-15 complex (IL-15C), which is IL-15 complexed with the IL-15Rα chain to extend *in vivo* half-life (65) mitigates CD8 T cell terminal exhaustion and promotes long-term efficacy of αPD-L1 or Dual therapy in the *KPC*2a model (23,66). However, the prior work focused on CD8 T cells, and both Th1 and Treg cells are theoretical targets of IL-15C. To determine the impact of IL-15C, cohorts of mice with orthotopic *KPC*2a tumors received αCD40 +αPD-L1 (Dual) or αCD40 + αPD-L1 + IL-15C (Triple) (**Figure 7I**) as described (23). We analyzed CD4 T cells using the MHC-II affinity enhanced CB54:I-A^b^ tetramer on day 14, because, by day 21, tumors are eradicated in 100% of the animals treated with Triple therapy (23), and tumor sizes were not different between the two cohorts (**Figure 7J**) indicating tumor mass was not a contributing factor for qualitative changes in CD4 T cells. The addition of IL-15C promoted an increase in frequency and number of tetramer-binding CD4 T cells in spleen and tumor (**Figure 7K**). IL-15C promoted tumor-specific T-bet^+^ Th1 cells, particularly in secondary lymphoid organs, while not expanding Treg cells (**Figure 7L, Supplementary Figure 8J-K**). Further, IL-15C significantly reduced PD-1 on tetramer^+^ CD4 T cells in all tissues (**Figure 7M**). Cxcr5, a Tfh marker, was drastically reduced in tetramer^+^ T cells in dLNs following the triple therapy (**Figure 7M, Supplementary Figure 8L**), suggesting that the addition of IL-15C biases differentiating tumor-specific CD4 T cells toward a Th1 state at the expense of Tfh. Further, the addition of IL-15C promoted Cxcr6^+^ and CD49a^+^Cxcr6^+^ tumor-specific CD4 T cells (**Figure 7N**), suggesting IL-15C with Dual promotes T cell residency traits and potentially enhanced trafficking to the tumor in response to Dual-induced *Cxcl16* produced by intratumoral myeloid cells (Fig 5). Lastly, we measured the function of intratumoral CD4 T cells by *ex vivo* restimulation. The addition of IL-15C decreased IL-10 production while increasing IFN-γ and TNF production (**Figure 7O**). Altogether, IL-15C augments Dual therapy by expanding antigen-specific Th1 cells without promoting Treg cells and enhances polyfunctional responses. Collectively, the above results support the critical role of tumor-specific CD4 T cells in PDA, and how to therapeutically enhance their antitumor activity with IL-15C-supported immunotherapy.

## DISCUSSION

Studies from our lab (6,10,23) and others (45,67,68) are uncovering a critical role for CD4 T helper cells in orchestrating a productive antitumor response in PDA. Mechanistic insight into tumor-specific CD4 T cells often derives from adoptive transfer of large numbers of monoclonal TCR-transgenic T cells with a fixed affinity and specific to standard model foreign antigens (9,54,69,70). Based on our discovery that CBR is immunogenic in B6 mice (22), we identified two novel I-A^b^-restricted CBR epitopes that are naturally processed and presented *in vivo*. Vaccination with MHC-II-restricted CBR epitopes generated protective immunity whereas vaccination with an immunodominant MHC-I epitope conferred no such protection. These data are consistent with prior study that vaccination with a single MHC-II restricted neoantigen conferred significant survival benefit in a melanoma model (8).

Identifying such antigen-specific CD4 T cells among the vast pool of polyclonal T cells is challenging due to their extremely low abundance. As such, we developed an MHC-II affinity-enhanced tetramer specific to CB54:I-Ab to track endogenous polyclonal tumor-specific CD4 T cells that naturally occur at a physiologic abundance. Indeed, we detected ∼100 CB54:I-A^b^-specific precursor CD4 T cells per mouse prior to antigen exposure. After tumor exposure, whereas CD8 T cells specific to CB_101-109_:H-2D^b^ can comprise over 20-50% of the CD8 T cell infiltrate (6,22,23), CB54:I-A^b^-specific CD4 T cells comprised a mere 2-7% of intratumoral CD4 T cells. Similarly, CD4 T cells specific to a naturally occurring mutant ITGB1 neoepitope found in a sarcoma model comprise ∼5% of the CD4 T cell infiltrate and were required for responses to immunotherapy (7). Thus, the CB54 model antigen produces a tumor-specific CD4 T cell population that is tractable and with an abundance comparable to CD4 T cells reactive to a naturally occurring neoantigen.

Using our CD4-affinity-enhanced MHC-II tetramer to capture the natural repertoire as it is shaped by tumor progression and immunotherapy, we demonstrate PDA-specific CD4 T cells are primed in *KPC*2a-bearing mice, harbor distinct phenotypic states in dLN and tumor. PDA-specific CD4 T cells in the dLN were biased to a Tfh-like state while in tumors they upregulate Cxcr6, supporting differentiation in this non-lymphoid site. Recent studies have identified a population of stem-like CD4 T cells that share key developmental and transcriptional features with Tfh cells, including expression of PD-1, TCF-1, BCL6, CXCR5, and IL-21 and are an implicated reservoir of CD4 T cells during chronic antigen encounter (35,71–75). PD-1 signaling has been shown to impact Tfh positioning and function (76,77). Following PD-L1 blockade, clonally expanded Tfh cells were identified intratumorally, by scRNA-seq, and tumor-specific Tfh expanded dramatically in dLNs. These data suggest that the progenitor CD4 T cell that responds to PD-1/PD-L1 blockade include Tfh cells. Critically, tumor-specific CD4 T cells infiltrating PDA contract over time, and remaining cells co-expressed PD-1 and Lag3, consistent with chronic antigen encounter and co-inhibitory signaling. Following PD-L1 blockade, clonally expanded intratumoral Tfh-like cells were identified, suggesting the progenitor CD4 T cell that responds to PD-1/PD-L1 blockade may be among Tfh-like cells. This hypothesis is supported by the reported plasticity of the Tfh subset (30,75).

We show αCD40-mediated therapeutic effect is highly dependent on CD4 T cells during the effector phase, indicating αCD40 cannot compensate for a lack of CD4 T cell help. Agonistic αCD40 was the major driver of intratumoral CD4 T cell clonal expansion during combination therapies. These data are consistent with CD44⁺CD4 Th1 cells correlating with improved disease free survival following αCD40 in PDA patients (78). Given that *KPC* cells fail to upregulate MHC-II, CD4 T cells must be acting indirectly (10). Such indirect antitumor effects of CD4 T cells may be through programming antitumor myeloid cells that are toxic to tumor cells and/or tumor-associated vasculature (54,79,80). We speculate, the CD4 T cell-dependent mechanism is driven by IFN-γ receptor signaling on host cells as we previously showed that efficacy of Dual therapy was dependent on Ifngr1 and independent of Tnfr1 (10,11).

Through single cell transcriptional analyses, we identify that αCD40 alters intratumoral myeloid cell composition, consistent with earlier work that αCD40 can activate monocytes with tumoricidal activity (81). αCD40 drove *Stat1*⁺ proinflammatory TAMs with genes involved in antigen presentation and T cell recruitment. Indeed, this *Stat1*^+^ population was highly enriched for T cell attracting chemokines *Cxcl9, Cxcl16*, and *Ccl5*, that bind Cxcr3, Cxcr6, and Ccr5. A comparable activation of the myeloid compartment has been reported in human trials in immune checkpoint blockade in head and neck squamous cell carcinoma (82), suggesting that some of the effects of αCD40 on myeloid cells are dependent on T cell activation. Indeed, CD4 T cells were required for intratumoral myeloid cell acquisition of tumor antigen. Effects of Dual therapy on CD4 T cell activation, tumor antigen uptake, and myeloid transcriptional changes likely facilitate the observed intratumoral triad formation.

MHC-II on Ccr2^+^ myeloid and myeloid-derived cells was critical for the antitumor effects of αCD40, TLS formation, and ICOS upregulation on tumor-specific CD4 T cells. ICOS is a costimulatory protein induced by TCR signaling on CD4 T cells (83) and ICOS:ICOSL signaling induces lymphotoxin a (LTα3) that promotes chemokine production for lymphocyte recruitment and TLS assembly (58). TLS have been identified in numerous human solid tumors, including PDA and often correlate with improved outcomes (17,84,85). However, TLS are not typically detected in murine tumor models, presumably because the reliance on implantable tumor cell lines that rapidly outpace the formation of immune aggregates. Our finding that αCD40 promotes TLS formation in orthotopic PDA is consistent with a study that showed αCD40 stimulation of B cells promotes TLS in the brain of glioma-bearing mice (86), and provides a way to study their formation going forward. The fact that B cells played no detectable role during immunotherapy here, can be explained by low B cell abundance in tumors prior to therapy and that B cells lack the same phagocytic capacity as myeloid cells.

To interrogate the clinical relevance of our findings, we assessed CD4 T cells in human HNSCC neoadjuvant immunotherapy cohorts (60). Across distinct immune checkpoint blockade regimens, clinical response was accompanied by shifts in the CD4 T cell compartment including promotion of a Th1 effector cells and a concurrent destabilization of Treg cells. Similar alterations in the Th1:Treg ratio were unleased through CTLA-4 blockade in a metastatic PDA setting (6). Agonistic αCD40 can promote the conversion of Treg cells to “ExTreg” Th1-like cells in pancreatic tumors 2 days after administration (45). Here, we show tumor-specific Treg cells are decreased in tumors 1 week after single αCD40 treatment. However, by 2 weeks post treatment, tumor-specific Tregs markedly rebound, to an extent that is even beyond control mice. As PD-L1 blockade interfered with this rebound, our data suggest PD-1:PD-L1 signaling may promote conversion and may explain why PD-L1 blockade has synergistic effects when combined with αCD40 (11,12). As these changes occurred across different immune checkpoint blockade regimens and between mouse and human solid tumors, Th1 potentiation coupled to Treg restraint may be generalizable determinant of immunotherapy response.

IL-2Rβ is a shared receptor subunit between IL-2 and IL-15 cytokines. Consistent with known expression pattern of IL-2Rβ, Th1 cells infiltrating PDA expressed abundant *Il2rb*. Cells enriched for *Il2rb* were also enriched for *Tbx21,Cxcr6* and *Foxp3*, and inversely with *Tcf7* across both mouse and human tumor-infiltrating CD4 T cells. IL-15C can promote Treg cell expansion that can be regulated by IFN-γ in diabetes model (87). Thus, the context of IL-15C therapy is critical. When combined with Dual therapy that promotes IFN-γ production by antigen-specific T cells (11), IL-15C biases toward Th1 cell expansion in tumors and secondary lymphoid organs. Functionally, IL-15C delivered with Dual therapy decreased IL-10 production while increasing IFNγ by tumor-infiltrating CD4 T cells. Thus, IL-15C in with Dual therapy can can further enhance tumor-specific Th1 cells without promoting Treg cells and is a likely mechanistic underpinning the success of this combination therapy and potentially as well as a factor that mitigates CD8 terminal exhaustion in PDA (23).

## LIMITATIONS OF THE STUDY

We employed a model antigen for interrogation of tumor-specific CD4 T cell responses which may not reflect all relevant antitumor CD4 T cell responses in humans. Antigen-specific CD4 T cell proliferative burst is tightly regulated as small numbers of cells are required to mediate immunological effects (54,88). As a result, even with an MHC-II affinity enhanced tetramer, the number of antigen-specific T cells available for analysis was relatively low as compared to our studies with CD8 T cells.

## METHODS

### Animals

All animal studies were approved by the University of Minnesota Institutional Animal Care and Use Committee (IACUC). Six- to 12-week-old male and female C57BL/6J (#000664) mice were purchased from The Jackson Laboratory. H2-Ab1fl/fl (#013181) mice were crossed with CCR2-CreERT2-GFP (#035229) mice to generate CCR2-CreERT2-GFP × H2-Ab1fl/fl (CCR2^ΔMHCII^) experimental and H2-Ab1fl/fl (CCR2^WT^) littermate control mice. For B cell-specific MHC-II ablation, H2-Ab1fl/fl mice were crossed with Cd19-Cre ( #006785, The Jackson Laboratory) to generate Cd19-Cre × H2-Ab1fl/fl and H2-Ab1fl/fl control mice. All strains were genotyped using in-house or Transnetyx protocols. Animals were maintained in specific pathogen-free conditions at the University of Minnesota Research Animal Resources facility with free access to food and water on a 12-hour light-dark cycle.

### Tumor Cell Lines

KPC2 primary pancreatic ductal tumor epithelial cells were derived from a C57BL/6J *Kras*^LSL-G12D/+^; *Trp53*^LSL-R172H/+;^ *p48*-Cre (*KPC*) mouse with invasive PDA. *KPC*2a cells were generated by retroviral transduction of *KPC*2 cells with click beetle red luciferase (CB) linked to eGFP (CBR-eGFP) and sorting for eGFP^+^ cells as described (22). Tumor cells were cultured in DMEM/F12 (Gibco) supplemented with 10% FBS (Gibco), 2.5 mg/mL amphotericin B (Gibco), 100 mg/mL penicillin/streptomycin (Gibco), and 2.5 g/L dextrose (Fisher Scientific) at 37°C and 5% CO2. Cell lines were tested monthly for Mycoplasma and maintained below passage 15. Cells were passaged using 0.25% trypsin-EDTA (Thermo Fisher).

### I-A^b^ Peptide Binding Prediction

Candidate MHC-II-restricted 11-mer peptides in CB were identified using two complementary binding prediction approaches: NetMHCIIpan and a custom algorithm developed by the Marc Jenkins laboratory (University of Minnesota). Top candidates were ranked by predicted binding affinity to I-A^b^ and selected for functional validation.

### TriVax Peptide Vaccination

Mice were vaccinated intraperitoneally with 100 μg CB54-64 (EFFEATVLLAQ), CB440-450 (KYKGSQVAPAE), and/or CB101-109 (VAPVNESYI) peptide (Genscript) combined with 50 μg agonistic αCD40 (FGK45, BioXCell) and 50 μg Poly(I:C) (Biotechne) in sterile saline to a final volume of 200 μl as described (22)

### Enzyme-Linked ImmunoSpot (ELISpot) Assay

IFN-γ ELISpot assays were performed using a mouse IFN-γ ELISpot kit (Mabtech) according to the manufacturer’s instructions. Briefly, 96-well PVDF plates were coated overnight with anti-IFN-γ capture antibody. Splenocytes from KPC2a tumor-bearing mice or TriVax-vaccinated mice were isolated as described and plated at 2 × 10^5^ cells per well. Cells were stimulated with 10 μg/ml CB54-64, CB440-450, or anchor-substituted control peptides for 18-20 hours at 37°C and 5% CO_2_. Wells were washed and incubated with biotinylated detection antibody, followed by streptavidin-alkaline phosphatase and BCIP/NBT substrate development. Spots were enumerated using an ImmunoSpot automated reader (Cellular Technology Limited).

### Orthotopic Tumor Implantation

Orthotopic tumor implantation was performed similar as we described (22). Mice received 1 mg/kg slow-release buprenorphine subcutaneously for analgesia prior to surgery and anesthetized with continuous 2-5% isoflurane. Abdominal hair was removed with clippers and Nair (Church & Dwight), and the skin was sterilized with a series of alternating ethanol and 10% betadine washes. Upon reaching surgical plane anesthesia, a small incision was made in the left lateral abdomen followed by a small peritoneal incision to access the pancreas. A total of 1 x 10^5^ tumor cells in 20 μl of 60% Growth Factor Reduced Matrigel (Corning) were injected into the pancreatic parenchyma using an insulin syringe (Covidien). The peritoneum was closed with absorbable sutures (Ethicon) and skin was closed with 9 mm wound clips (Fine Science Tools). Mice were monitored daily for 5 days post-surgery and received carprofen (10 mg/kg, subcutaneous) for 3 days post-operatively.

### In Vivo Antibody Treatments

Antibodies were diluted to 200 µL in sterile saline and administered intraperitoneally (i.p.) with a 30G needle. For immunotherapy studies, mice received a single dose of 100 μg agonistic αCD40 (FGK45, BioXCell) on day 7, alone or in combination with 200 μg αPD-L1 (10F.9G2, BioXCell) on days 7, 10, and 12 post-tumor implantation. For triple therapy, IL-15 complex (IL-15C) was prepared by combining 7 μg IL-15Rα-Fc (Biotechne) with 0.75 μg IL-15 (BioLegend) in sterile saline, incubated at 37°C for 30 minutes, and administered i.p. on days 7 and 9 post-tumor implantation as described (23,66).

### In vivo depletions

For CD4 T cell depletion, mice received 200 μg anti-CD4 (GK1.5, BioXCell) i.p. on days −1, 2, 6, and 10 relative to tumor implantation. For CD4 T cell depletion in Dual therapy treated mice, 200 μg anti-CD4 was administered on days 6 and 10 post tumor implantation. For B cell depletion, mice received 400 μg anti-CD20 (SA271G2) i.p. on day −1 relative to tumor implantation. Depletion efficiency was confirmed by flow cytometry.

### Preparation of Single-Cell Suspensions

Spleens and pancreatic draining lymph nodes were mechanically dissociated over a 40 μm cell strainer. Splenocyte suspensions were subjected to ACK red blood cell lysis (Gibco) for 3 minutes at room temperature, quenched with complete media (DMEM + 2% FBS + 100 μg/mL penicillin/streptomycin + 2 mM L-glutamine), and centrifuged at 350 × g for 5 minutes at 4°C. Tumors were dissected from adjacent normal pancreas, weighed, minced, and digested in 0.5 mg/mL Collagenase IV (Sigma Aldrich) in DMEM for 15 minutes at 37°C with vigorous shaking (180 rpm), followed by mechanical dissociation over a 70 μm strainer and two wash steps to remove cell debris and pancreatic enzymes. All single-cell suspensions were maintained on ice in complete media prior to staining.

### MHC-II Tetramer Generation

Affinity-enhanced I-Ab (I-Ab-4E) monomers bearing CB54-64 (EFFEATVLLAQ) were generated and biotinylated as previously described (16). Tetramers were assembled by mixing biotinylated monomer with PE- or APC-conjugated streptavidin at a 4:1 molar ratio.

### Flow cytometry

Tumors were minced and digested with Collagenase IV (Sigma Aldrich) at 37°C for 15 minutes with vigorous shaking (180rpm), then mechanically dissociated to single-cell suspensions and washed twice to remove cell debris. Cells were stained antibodies diluted 1:200 in FACS buffer (PBS + 2.5% FBS) with 1:100 Fc block (CD16/32, Tonbo) for 45 minutes at 4°C in the dark. Zombie NIR (Tonbo) was included at 1:500 to exclude dead cells. For MHC-II tetramer staining, cells were incubated with two CB54:I-A^b^ tetramers conjugated to different fluorophores (1:100) and surface staining antibodies with gentle shaking (120 rpm) for 45 minutes at room temperature. For intracellular transcription factor staining, cells were fixed and permeabilized using the Foxp3 fixation/permeabilization kit (Tonbo) and stained for 30 minutes at room temperature with gentle shaking (120rpm) with antibodies diluted in 1X permeabilization buffer. For intracellular cytokine staining (ICS), single-cell suspensions were stimulated for 4.5 hours at 37°C with 10 µg/mL CB54-64 peptide (Genscript) or Cell Stimulation Cocktail (PMA/ionomycin, 1:500, Tonbo) in the presence of GolgiPlug (1:1000, BD) and GolgiStop (1:1500, BD). Following surface staining, cells were fixed and permeabilized using the BD Cytofix/Cytoperm kit and stained with secreted factor-targeting antibodies for 1 hour at 4°C. For tumor antigen internalization assays, unfixed single-cell suspensions from GFP-expressing *KPC*2a-bearing tumors were stained for myeloid surface markers and acquired the same day to quantify GFP fractions among TAMs and DCs. Cell counting beads (Sigma Aldrich, 100 µL/sample) were added immediately prior to acquisition for absolute cell number calculations. If cells were not yet fixed (lacked intracellular staining), then they were fixed in 2% PFA or fixation buffer (Tonbo) for 10-15 minutes at room temperature in the dark prior to acquisition. Cells were acquired within 24 hours on a Cytek Aurora spectral flow cytometer (SpectroFlo software). Data were analyzed using FlowJo v10 (BD).

### High-resolution ultrasound

Tumor-bearing mice were anesthetized with continuous 2-4% isoflurane and abdominal hair was removed with clippers and Nair. Tumors were identified by anatomic landmarks, hypoechoic appearance, and defined margins using a Vevo F2 high-resolution ultrasound system (Visual Sonics). Tumor volume was calculated using the modified ellipsoidal formula: 0.5 x (length x width^2^).

### Immunofluorescence Microscopy

Tissues were embedded in OCT (Tissue-Tek), snap-frozen, and stored at −80°C. Cryosections (7 μm) were cut using a cryostat (Leica), mounted on glass slides, and fixed in ice-cold acetone for 10 minutes at −20°C. Sections were rehydrated with PBS + 1% BSA, blocked with 5% BSA/PBS for 1 hour at room temperature in a humidified chamber, and incubated with primary antibodies against CD4, CD8α, F4/80, CD11c, CD19, or pan-cytokeratin diluted in 1% BSA/PBS for 1 hour at room temperature. Slides were washed three times in PBS and mounted with ProLong Gold Antifade with DAPI (Invitrogen). Images were acquired on a Thunder Imager widefield microscope platform (Leica). Quantification of tertiary lymphoid structures (TLS), immune cell aggregates, and intratumoral immune triads (CD4^+^: CD8^+^: CD11c^+^ cells in <20 μm proximity) was performed using QuPath software (v0.7.0). TLS were defined by dense cell nuclei aggregates infiltrated by both CD4 T cells and CD19^+^ B cells. Tumor core was manually annotated as regions of architecturally disordered cytokeratin (CK)^+^ regions which is characteristic of invasive carcinoma. Peritumor was defined as the region within 100 μm of the tumor core border, generated by first expanding the tumor core ROI by 100 μm using a custom Groovy script in QuPath followed by subtracting the tumor core ROI from the expanded ROI using a second custom Groovy script.

### Tamoxifen Administration

To induce Cre-mediated MHC-II deletion, 5 mg of tamoxifen (Sigma-Aldrich) resuspended in corn oil was delivered by oral gavage on days 3, 6, 9, and 12 post-tumor, as we described (10).

### In Vitro IFN-γ Stimulation of Tumor Cells

1 x10^6^ *KPC*2 or *KPC*2a tumor cells were seeded on 10 cm plates and allowed to adhere for 24 hours. Recombinant murine IFN-γ (Fisher) was added at 10 ng/mL. After 48 hours, cells were harvested with 0.25% trypsin-EDTA and analyzed for MHC class I (H2-K^b^, H2-D^b^) and MHC class II (I-A/I-E) surface expression by flow cytometry.

### Mouse single cell RNA-sequencing

We analyzed scRNA-seq from murine orthotopic *KPC*2 and *KPC*2a tumor-bearing mice (6,23) and are publicly available through Gene Expression Omnibus: GSE330483 and GSE285902. Sample demultiplexing was performed bioinformatically using hashtag oligo (HTO) counts. Preprocessing excluded genes detected in fewer than 10 cells, cells with fewer than 200 features, and cells with greater than 25% mitochondrial gene content. Data normalization and scaling were performed using NormalizeData and ScaleData in Seurat (v5.2.1). Variable features were identified with FindVariableFeatures, and batch correction across samples was performed with Harmony. CD4+ T cells were isolated by subsetting T cell clusters followed by filtering on Cd4 transcript expression above the non-zero 1st percentile, followed by reclustering and UMAP visualization. Myeloid cell subclustering was done by removing B cell (*Cd19*), T cell (*Trac*), and neutrophil (*Csf3r*) clusters, followed by reclustering and UMAP visualization. Differentially expressed genes (DEG) between cells in the control condition compared to the treated groups was conducted using Seurat’s FindMarkers function. For myeloid cell GSEA analysis, DEG ranked by p-value were analyzed against the GO “BP” ontology using clusterProfiler (org.Mm.eg.db, set sizes 15–500, Benjamini-Hochberg FDR < 0.05). Visualization used ggplot2 and pheatmap.

### Human single cell RNA-sequencing data analysis

Li et al. single cell RNA-seq and CITE-seq data (GEO: GSE288199, NCT04080804 (62)) were accessed using the GEOquery R package. Samples were imported and merged using Seurat, with chemistry version (10x Genomics 5’ v1 or v2) and sample identity included for downstream batch correction variables in Harmony integration. Quality control filtering was applied to remove low-quality cells with greater than 20% mitochondrial gene expression, and cells with fewer than 200 or greater than 10,000 detected features were excluded. Prior to dimensionality reduction and clustering, mitochondrial genes, ribosomal protein genes (RPL/RPS), and TCR variable region genes (TRAV/TRBV) were excluded from the set of highly variable features.

CD4 T cells were isolated by removing a cluster of myeloid contaminants (CD14^+^), filtering for CD4 RNA expression greater than zero, and filtering for CD4 CITE-seq antibody expression above the non-zero first percentile. Cross-patient samples underwent normalization via SCTransform followed by dimensionality reduction. Clustering was performed at resolution 0.17 yielding six transcriptionally defined CD4+ subpopulations used for downstream analyses. Gene signature module scores were calculated for each CD4+ T cell using the AddModuleScore() function in Seurat. Two published gene signatures were scored: a stem-like progenitor signature (BCL6, IL7R, CCR7, TCF7, BACH2, CXCR5, SELL) derived from tumor-infiltrating CD8+ T cells in human colorectal cancer (64), and a germinal center formation (GO:0002467). For subset-specific analyses, Treg cells were defined as cells with FOXP3 RNA expression greater than zero, and Th1 cells were defined as cells with a positive Th1 module score calculated from a published CellMarker2 signature (CL:0000545; undefined normal Th1). Pre-versus post-treatment differences in module scores were assessed per treatment arm using the Wilcoxon signed-rank test with BH correction applied across the three arm-wise comparisons per signature.

### NicheNet analysis

Intercellular ligand-target predictions were performed using NicheNet (59) on the KPC2a single-cell RNA-seq dataset, with CD4 T cells defined as the sender population and myeloid cells as the receiver population. The gene set of interest comprised differentially expressed genes between control and Dual therapy-treated myeloid cells, identified using Seurat’s FindMarkers, with all genes expressed in the receiver population used as the background. Ligands were prioritized by the area under the precision-recall curve (AUPR) between their predicted target genes and the observed differential expression signature, and top-ranked ligands were retained. Regulatory potential scores between prioritized ligands and their predicted target genes were visualized as a heatmap.

### Statistical Analysis

Statistical analyses were performed using GraphPad Prism (v10). Two-group comparisons used an unpaired, two-tailed Student’s t-test. Comparisons among >2 groups used one-way ANOVA with Tukey’s or Dunnett’s post-test. If normality testing failed, Mann-Whitney or Kruskal-Wallis non-parametric or lognormal tests were applied. Data are presented as mean ± SEM unless otherwise noted. P < 0.05 was considered statistically significant (*p<0.05, **p<0.01, ***p<0.001, ****p< 0.0001).

## Acknowledgements

We thank Brandon M. Larsen, Grant H. Hickok, Audrey L. Hilk, Alexander K. Tsai, Cara-lin Lonetree, and Ebony Miller for experimental support and Hezkiel A. Nanda for initial bioinformatics consultation. We thank Jesse Williams and Samuel Becker for their input on myeloid cell experiments and transcriptional data. We thank the University of Minnesota Genomics Core (RRID: SCR_012413), Imaging Centers (RRID: SCR_020997), the Center for Immunology Imaging Core, Flow Cytometry Core, and Research Animal Resource facility. We also thank the NIH Tetramer Core Facility (RRID:SCR_026557) for generating traditional (non-affinity-enhanced) CB54:I-A^b^ peptide/MHC tetramers used in initial studies.

## Author contributions

Conceptualization: E.C.H. and I.M.S. Formal Analysis: E.C.H., S.M.B., and I.M.S. Funding acquisition: I.M.S. Investigation: E.C.H., Z.C.S., S.M.B., A.L.B., M.A.E., R.P., and A.G. Methodology: S.A., D.T. Supervision: S.S., D.T., and I.M.S. Visualization: E.C.H., S.M.B., R.P., A.G., I.M.S. Writing—original draft: E.C.H. and I.M.S. Writing—review & editing: E.C.H., Z.C.S.,A.L.B., and I.M.S.

## Author Approval

all authors have seen and approved the manuscript, and that it hasn’t been accepted or published elsewhere.

## References

1. American Cancer Society. Cancer Facts & Figures 2023. Atlanta: American Cancer Society; 2023.

2. O’Reilly EM, Oh DY, Dhani N, Renouf DJ, Lee MA, Sun W, et al. Durvalumab with or Without Tremelimumab for Patients with Metastatic Pancreatic Ductal Adenocarcinoma: A Phase 2 Randomized Clinical Trial. JAMA Oncology. 2019;5:1431–8.

3. Schmiechen ZC, Stromnes IM. Mechanisms Governing Immunotherapy Resistance in Pancreatic Ductal Adenocarcinoma. Frontiers in Immunology. 2021;11.

4. Montauti E, Oh DY, Fong L. CD4+ T cells in antitumor immunity. Trends in Cancer. Elsevier; 2024;10:969– 85.

5. Speiser DE, Chijioke O, Schaeuble K, Münz C. CD4+ T cells in cancer. Nat Cancer. Nature Publishing Group; 2023;4:317–29.

6. Schmiechen ZC, Cruz-Hinojoza E, Hilk AL, Ellefson MA, Tsai AK, Dres OM, et al. Treg cells promote immunotherapy-induced immune evasion by restraining CD4 T cell control of MHC-I–deficient metastatic pancreatic cancer. Science Immunology. American Association for the Advancement of Science; 2026;11:eadz4302.

7. Alspach E, Lussier DM, Miceli AP, Kizhvatov I, DuPage M, Luoma AM, et al. MHC-II neoantigens shape tumour immunity and response to immunotherapy. Nature. Nature Publishing Group; 2019;574:696–701.

8. Kreiter S, Vormehr M, van de Roemer N, Diken M, Löwer M, Diekmann J, et al. Mutant MHC class II epitopes drive therapeutic immune responses to cancer. Nature. 2015;520:692–6.

9. Espinosa-Carrasco G, Chiu E, Scrivo A, Zumbo P, Dave A, Betel D, et al. Intratumoral immune triads are required for immunotherapy-mediated elimination of solid tumors. Cancer Cell. 2024;42:1202–1216.e8.

10. Patterson MT, Burrack AL, Xu Y, Hickok GH, Schmiechen ZC, Becker S, et al. Tumor-specific CD4 T cells instruct monocyte fate in pancreatic ductal adenocarcinoma. Cell Reports. Elsevier; 2023;42.

11. Burrack AL, Rollins MR, Spartz EJ, Mesojednik TD, Schmiechen ZC, Raynor JF, et al. CD40 Agonist Overcomes T Cell Exhaustion Induced by Chronic Myeloid Cell IL-27 Production in a Pancreatic Cancer Preclinical Model. J Immunol. Baltimore, Md.: 1950; 2021;206:1372–84.

12. Vonderheide RH. CD40 Agonist Antibodies in Cancer Immunotherapy. Annual Review of Medicine. Annual Reviews; 2020;71:47–58.

13. Padrón LJ, Maurer DM, O’Hara MH, O’Reilly EM, Wolff RA, Wainberg ZA, et al. Sotigalimab and/or nivolumab with chemotherapy in first-line metastatic pancreatic cancer: clinical and immunologic analyses from the randomized phase 2 PRINCE trial. Nat Med. Nature Publishing Group; 2022;28:1167–77.

14. O’Hara MH, O’reilly EM, Varadhachary G, Wolff RA, Wainberg ZA, Ko AH, et al. CD40 agonistic monoclonal antibody APX005M (sotigalimab) and chemotherapy, with or without nivolumab, for the treatment of metastatic pancreatic adenocarcinoma: an open-label, multicentre, phase 1b study. The Lancet Oncology. 2021;22:118–31.

15. Kotov DI, Jenkins MK. Peptide:MHCII Tetramer-Based Cell Enrichment for the Study of Epitope-Specific CD4+ T Cells. Curr Protoc Immunol. 2019;125:e75.

16. Dileepan T, Malhotra D, Kotov DI, Kolawole EM, Krueger PD, Evavold BD, et al. MHC class II tetramers engineered for enhanced binding to CD4 improve detection of antigen-specific T cells. Nat Biotechnol. 2021;39:943–8.

17. Stromnes IM, Hulbert A, Pierce RH, Greenberg PD, Hingorani SR. T-cell Localization, Activation, and Clonal Expansion in Human Pancreatic Ductal Adenocarcinoma. Cancer Immunol Res. 2017;5:978–91.

18. Bailey P, Chang DK, Nones K, Johns AL, Patch A-M, Gingras M-C, et al. Genomic analyses identify molecular subtypes of pancreatic cancer. Nature. Nature Publishing Group; 2016;531:47–52.

19. Balli D, Rech AJ, Stanger BZ, Vonderheide RH. Immune Cytolytic Activity Stratifies Molecular Subsets of Human Pancreatic Cancer. Clin Cancer Res. 2017;23:3129–38.

20. Hingorani SR, Petricoin EF, Maitra A, Rajapakse V, King C, Jacobetz MA, et al. Preinvasive and invasive ductal pancreatic cancer and its early detection in the mouse. Cancer Cell. 2003;4:437–50.

21. Hingorani SR, Wang L, Multani AS, Combs C, Deramaudt TB, Hruban RH, et al. Trp53R172H and KrasG12D cooperate to promote chromosomal instability and widely metastatic pancreatic ductal adenocarcinoma in mice. Cancer Cell. 2005;7:469–83.

22. Burrack AL, Spartz EJ, Raynor JF, Wang I, Olson M, Stromnes IM. Combination PD-1 and PD-L1 Blockade Promotes Durable Neoantigen-Specific T Cell-Mediated Immunity in Pancreatic Ductal Adenocarcinoma. Cell Rep. 2019;28:2140–2155.e6.

23. Schmiechen ZC, Nanda HA, Burrack AL, Hickok GH, Butler JZ, Cruz-Hinojoza E, et al. IL-15 Complex Enhances Agonistic Anti-CD40 + Anti-PDL1 by Correcting the T-bet to Tox Ratio in CD8+ T cells Infiltrating Pancreatic Ductal Adenocarcinoma. Cancer Immunol Res. 2025;13:847–66.

24. Burrack AL, Schmiechen ZC, Patterson MT, Miller EA, Spartz EJ, Rollins MR, et al. Distinct myeloid antigen-presenting cells dictate differential fates of tumor-specific CD8+ T cells in pancreatic cancer. JCI Insight. 2022;7:e151593.

25. Moon JJ, Chu HH, Pepper M, McSorley SJ, Jameson SC, Kedl RM, et al. Naive CD4(+) T cell frequency varies for different epitopes and predicts repertoire diversity and response magnitude. Immunity. 2007;27:203–13.

26. Gesbert F, Moreau J-L, Thèze J. IL-2 responsiveness of CD4 and CD8 lymphocytes: further investigations with human IL-2Rbeta transgenic mice. Int Immunol. 2005;17:1093–102.

27. Foulds KE, Zenewicz LA, Shedlock DJ, Jiang J, Troy AE, Shen H. Cutting edge: CD4 and CD8 T cells are intrinsically different in their proliferative responses. J Immunol. Baltimore, Md.: 1950; 2002;168:1528–32.

28. De Boer RJ, Homann D, Perelson AS. Different dynamics of CD4+ and CD8+ T cell responses during and after acute lymphocytic choriomeningitis virus infection. J Immunol. Baltimore, Md.: 1950; 2003;171:3928–35.

29. Weber JP, Fuhrmann F, Feist RK, Lahmann A, Al Baz MS, Gentz L-J, et al. ICOS maintains the T follicular helper cell phenotype by down-regulating Krüppel-like factor 2. J Exp Med. 2015;212:217–33.

30. Crotty S. Follicular helper CD4 T cells (TFH). Annu Rev Immunol. 2011;29:621–63.

31. Mabrouk N, Tran T, Sam I, Pourmir I, Gruel N, Granier C, et al. CXCR6 expressing T cells: Functions and role in the control of tumors. Front Immunol. 2022;13:1022136.

32. Hong S-W, Krueger PD, Osum KC, Dileepan T, Herman A, Mueller DL, et al. Immune tolerance of food is mediated by layers of CD4+ T cell dysfunction. Nature. 2022;607:762–8.

33. Metzger TC, Long H, Potluri S, Pertel T, Bailey-Bucktrout SL, Lin JC, et al. ICOS Promotes the Function of CD4+ Effector T Cells during Anti-OX40-Mediated Tumor Rejection. Cancer Res. 2016;76:3684–9.

34. Brennan M, DeBruin D, Nwokolo C, Hunt KS, Piening A, Donlin MJ, et al. T-Cell Expression of CXCL13 is Associated with Immunotherapy Response in a Sex-Dependent Manner in Patients with Lung Cancer. Cancer Immunol Res. 2024;12:956–63.

35. Cardenas MA, Prokhnevska N, Sobierajska E, Gregorova P, Medina CB, Valanparambil RM, et al. Differentiation fate of a stem-like CD4 T cell controls immunity to cancer. Nature. 2024;636:224–32.

36. Xu W, Zhao X, Wang X, Feng H, Gou M, Jin W, et al. The Transcription Factor Tox2 Drives T Follicular Helper Cell Development via Regulating Chromatin Accessibility. Immunity. 2019;51:826–839.e5.

37. Whiteside SK, Grant FM, Gyori DS, Conti AG, Imianowski CJ, Kuo P, et al. CCR8 marks highly suppressive Treg cells within tumours but is dispensable for their accumulation and suppressive function. Immunology. 2021;163:512–20.

38. Kim H-J, Barnitz RA, Kreslavsky T, Brown FD, Moffett H, Lemieux ME, et al. Stable inhibitory activity of regulatory T cells requires the transcription factor Helios. Science. New York, N.Y.; 2015;350:334–9.

39. Galpin KJC, Rodriguez GM, Maranda V, Cook DP, Macdonald E, Murshed H, et al. FGL2 promotes tumour growth and attenuates infiltration of activated immune cells in melanoma and ovarian cancer models. Sci Rep. 2024;14:787.

40. Yan J, Zhao Q, Gabrusiewicz K, Kong L-Y, Xia X, Wang J, et al. FGL2 promotes tumor progression in the CNS by suppressing CD103+ dendritic cell differentiation. Nat Commun. 2019;10:448.

41. Kidani Y, Nogami W, Yasumizu Y, Kawashima A, Tanaka A, Sonoda Y, et al. CCR8-targeted specific depletion of clonally expanded Treg cells in tumor tissues evokes potent tumor immunity with long-lasting memory. Proc Natl Acad Sci U S A. 2022;119:e2114282119.

42. Nettersheim FS, Brunel S, Sinkovits RS, Armstrong SS, Roy P, Billitti M, et al. PD-1 and CD73 on naive CD4+ T cells synergistically limit responses to self. Nat Immunol. 2025;26:105–15.

43. He L, Gu W, Wang M, Chang X, Sun X, Zhang Y, et al. Extracellular matrix protein 1 promotes follicular helper T cell differentiation and antibody production. Proc Natl Acad Sci U S A. 2018;115:8621–6.

44. Koch MA, Tucker-Heard G, Perdue NR, Killebrew JR, Urdahl KB, Campbell DJ. The transcription factor T-bet controls regulatory T cell homeostasis and function during type 1 inflammation. Nat Immunol. 2009;10:595–602.

45. Maltez VI, Arora C, Gribbin KP, Caruso B, Haerr ME, Sor R, et al. Agonistic anti-CD40 antibody treatment converts resident regulatory T cells into activated type 1 effectors within the tumor microenvironment. Immunity. 2026;59:1058–1074.e7.

46. Schumacher TN, Thommen DS. Tertiary lymphoid structures in cancer. Science. New York, N.Y.; 2022;375:eabf9419.

47. Rastogi I, Jeon D, Moseman JE, Muralidhar A, Potluri HK, McNeel DG. Role of B cells as antigen presenting cells. Front Immunol. 2022;13:954936.

48. Teillaud J-L, Houel A, Panouillot M, Riffard C, Dieu-Nosjean M-C. Tertiary lymphoid structures in anticancer immunity. Nat Rev Cancer. 2024;24:629–46.

49. Lehmann J, Thelen M, Kreer C, Schran S, Garcia-Marquez MA, Cisic I, et al. Tertiary Lymphoid Structures in Pancreatic Cancer are Structurally Homologous, Share Gene Expression Patterns and B-cell Clones with Secondary Lymphoid Organs, but Show Increased T-cell Activation. Cancer Immunol Res. 2025;13:323–36.

50. Damle SR, Carter JA, Goodsell KE, Pineda JMB, Dickerson LK, Jiang X, et al. Intratumoral Three-Cell-Type Clusters Are a Conserved Feature of Endogenous Antitumor Immunity. Cancer Immunology Research. 2026;14:205–18.

51. Baek M, DiMaio F, Anishchenko I, Dauparas J, Ovchinnikov S, Lee GR, et al. Accurate prediction of protein structures and interactions using a three-track neural network. Science. New York, N.Y.; 2021;373:871–6.

52. Maier B, Leader AM, Chen ST, Tung N, Chang C, LeBerichel J, et al. A conserved dendritic-cell regulatory program limits antitumour immunity. Nature. 2020;580:257–62.

53. DeNardo DG, Barreto JB, Andreu P, Vasquez L, Tawfik D, Kolhatkar N, et al. CD4+ T Cells Regulate Pulmonary Metastasis of Mammary Carcinomas by Enhancing Protumor Properties of Macrophages. Cancer Cell. Elsevier; 2009;16:91–102.

54. Kruse B, Buzzai AC, Shridhar N, Braun AD, Gellert S, Knauth K, et al. CD4+ T cell-induced inflammatory cell death controls immune-evasive tumours. Nature. 2023;618:1033–40.

55. Wiesolek HL, Bui TM, Lee JJ, Dalal P, Finkielsztein A, Batra A, et al. Intercellular Adhesion Molecule 1 Functions as an Efferocytosis Receptor in Inflammatory Macrophages. Am J Pathol. 2020;190:874–85.

56. Denda-Nagai K, Aida S, Saba K, Suzuki K, Moriyama S, Oo-Puthinan S, et al. Distribution and function of macrophage galactose-type C-type lectin 2 (MGL2/CD301b): efficient uptake and presentation of glycosylated antigens by dendritic cells. J Biol Chem. 2010;285:19193–204.

57. Browaeys R, Saelens W, Saeys Y. NicheNet: modeling intercellular communication by linking ligands to target genes. Nat Methods. 2020;17:159–62.

58. Nayar S, Pontarini E, Campos J, Berardicurti O, Smith CG, Asam S, et al. Immunofibroblasts regulate LTα3 expression in tertiary lymphoid structures in a pathway dependent on ICOS/ICOSL interaction. Commun Biol. 2022;5:413.

59. Yang Y, Yang X, Wang Y, Xu J, Shen H, Gou H, et al. Combined Consideration of Tumor-Associated Immune Cell Density and Immune Checkpoint Expression in the Peritumoral Microenvironment for Prognostic Stratification of Non-Small-Cell Lung Cancer Patients. Front Immunol. 2022;13:811007.

60. Li H, Zandberg DP, Kulkarni A, Chiosea SI, Santos PM, Isett BR, et al. Distinct CD8+ T cell dynamics associate with response to neoadjuvant cancer immunotherapies. Cancer Cell. 2025;43:757–775.e8.

61. Doan AE, Mueller KP, Chen AY, Rouin GT, Chen Y, Daniel B, et al. FOXO1 is a master regulator of memory programming in CAR T cells. Nature. 2024;629:211–8.

62. Liu J, Zeng J, Zhou T, Wu M, Weng X. Increased CD103-CD8+ TILs with TPEX phenotype replenish anti-tumor T cell pool in mismatch repair-proficient CRC. Cancer Immunol Immunother. 2025;74:358.

63. Cao Z, Wara AK, Icli B, Sun X, Packard RRS, Esen F, et al. Kruppel-like factor KLF10 targets transforming growth factor-beta1 to regulate CD4(+)CD25(-) T cells and T regulatory cells. J Biol Chem. 2009;284:24914–24.

64. Chen X, Wu X, Zhou Q, Howard OMZ, Netea MG, Oppenheim JJ. TNFR2 is critical for the stabilization of the CD4+Foxp3+ regulatory T. cell phenotype in the inflammatory environment. J Immunol. Baltimore, Md.: 1950; 2013;190:1076–84.

65. Mortier E, Quéméner A, Vusio P, Lorenzen I, Boublik Y, Grötzinger J, et al. Soluble Interleukin-15 Receptor α (IL-15Rα)-sushi as a Selective and Potent Agonist of IL-15 Action through IL-15Rβ/γ. Journal of Biological Chemistry. 2006;281:1612–9.

66. Burrack AL, Tsai AK, Ellefson MA, Schmiechen ZC, Larsen BM, Burrack KS, et al. IL-15 complex enhances therapeutic efficacy of anti-PD-L1 in a T cell-dependent and NK cell-independent manner in a murine model of pancreatic ductal adenocarcinoma. J Immunol. Baltimore, Md.: 1950; 2025;214:3228– 37.

67. Wattenberg MM, Coho H, Herrera VM, Graham K, Stone ML, Xue Y, et al. Cancer immunotherapy via synergistic coactivation of myeloid receptors CD40 and Dectin-1. Sci Immunol. 2023;8:eadj5097.

68. Huffman AP, Lin JH, Kim SI, Byrne KT, Vonderheide RH. CCL5 mediates CD40-driven CD4+ T cell tumor infiltration and immunity. JCI Insight. 2020;5:e137263, 137263.

69. Brightman SE, Becker A, Thota RR, Naradikian MS, Chihab L, Zavala KS, et al. Neoantigen-specific stem cell memory-like CD4+ T cells mediate CD8+ T cell-dependent immunotherapy of MHC class II-negative solid tumors. Nat Immunol. 2023;24:1345–57.

70. Wolf SP, Leisegang M, Steiner M, Wallace V, Kiyotani K, Hu Y, et al. CD4+ T cells with convergent TCR recombination reprogram stroma and halt tumor progression in adoptive therapy. Sci Immunol. 2024;9:eadp6529.

71. Wen L, Su C-H, Potemkin N, Dryburgh L, Qin L, Li S, et al. Stem-like precursors of exhausted Th cells upheld by a Tox-Myb-Eomes transcriptional hierarchy propagate Th cell responses in chronic infection. Immunity. 2026;S1074–7613(26)00231-1.

72. Xia Y, Sandor K, Pai JA, Daniel B, Raju S, Wu R, et al. BCL6-dependent TCF-1+ progenitor cells maintain effector and helper CD4+ T cell responses to persistent antigen. Immunity. 2022;55:1200–1215.e6.

73. Zou D, Yin Z, Yi SG, Wang G, Guo Y, Xiao X, et al. CD4+ T cell immunity is dependent on an intrinsic stem-like program. Nat Immunol. 2024;25:66–76.

74. Sato Y, Jain A, Ohtsuki S, Okuyama H, Sturmlechner I, Takashima Y, et al. Stem-like CD4+ T cells in perivascular tertiary lymphoid structures sustain autoimmune vasculitis. Sci Transl Med. 2023;15:eadh0380.

75. Künzli M, Schreiner D, Pereboom TC, Swarnalekha N, Litzler LC, Lötscher J, et al. Long-lived T follicular helper cells retain plasticity and help sustain humoral immunity. Science Immunology. American Association for the Advancement of Science; 2020;5:eaay5552.

76. Ruggiu M, Guérin MV, Corre B, Bardou M, Alonso R, Russo E, et al. Anti-PD-1 therapy triggers Tfh cell-dependent IL-4 release to boost CD8 T cell responses in tumor-draining lymph nodes. J Exp Med. 2024;221:e20232104.

77. Shi J, Hou S, Fang Q, Liu X, Liu X, Qi H. PD-1 Controls Follicular T Helper Cell Positioning and Function. Immunity. 2018;49:264–274.e4.

78. Blise KE, Sivagnanam S, Betts CB, Betre K, Kirchberger N, Tate BJ, et al. Machine Learning Links T-cell Function and Spatial Localization to Neoadjuvant Immunotherapy and Clinical Outcome in Pancreatic Cancer. Cancer Immunol Res. 2024;12:544–58.

79. Lian Q, Nie J, Singh J, Chen Q, Matta J, Chan W, et al. CD4+ T cells impair tumor growth through IL-3 and TNF-dependent vascular damage. Science. New York, N.Y.; 2026;392:eads7910.

80. Kim SI, Haerr ME, Al-Ghezi M, Chen C, Zhang Y, Phipps JL, et al. CD4+ T Cells Mediate MHC-Deficient Tumor Rejection and Endothelial Cell Reprogramming. Cancer Immunol Res. 2026;14:107–21.

81. Beatty GL, Chiorean EG, Fishman MP, Saboury B, Teitelbaum UR, Sun W, et al. CD40 agonists alter tumor stroma and show efficacy against pancreatic carcinoma in mice and humans. Science. New York, N.Y.; 2011;331:1612–6.

82. Franken A, Bila M, Mechels A, Kint S, Van Dessel J, Pomella V, et al. CD4+ T cell activation distinguishes response to anti-PD-L1+anti-CTLA4 therapy from anti-PD-L1 monotherapy. Immunity. 2024;57:541–558.e7.

83. Hutloff A, Dittrich AM, Beier KC, Eljaschewitsch B, Kraft R, Anagnostopoulos I, et al. ICOS is an inducible T-cell co-stimulator structurally and functionally related to CD28. Nature. 1999;397:263–6.

84. Zou X, Lin X, Cheng H, Chen Y, Wang R, Ma M, et al. Characterization of intratumoral tertiary lymphoid structures in pancreatic ductal adenocarcinoma: cellular properties and prognostic significance. J Immunother Cancer. 2023;11:e006698.

85. Amisaki M, Zebboudj A, Yano H, Zhang SL, Payne G, Chandra AK, et al. IL-33-activated ILC2s induce tertiary lymphoid structures in pancreatic cancer. Nature. 2025;638:1076–84.

86. van Hooren L, Vaccaro A, Ramachandran M, Vazaios K, Libard S, van de Walle T, et al. Agonistic CD40 therapy induces tertiary lymphoid structures but impairs responses to checkpoint blockade in glioma. Nat Commun. 2021;12:4127.

87. Xia J, Liu W, Hu B, Tian Z, Yang Y. IL-15 promotes regulatory T cell function and protects against diabetes development in NK-depleted NOD mice. Clinical Immunology. 2010;134:130–9.

88. Künzli M, Masopust D. CD4+ T cell memory. Nat Immunol. 2023;24:903–14.

