## Supplementary Figures 1-8 for "Tumor-specific CD4 T cells cooperate with myeloid cells to remodel the pancreatic tumor microenvironment and enable effective immunotherapy"

**Cruz-Hinojoza et al. Supplementary Figures 1- 8**

#### Supplementary Figure 1

A

| Peptide | Undecamer Sequence | Jenkins score | Nonamer Core | NetMHC-II 2.1 Predicted Affinity(nM) |
|---|---|---|---|---|
| CB440-450 | KYKGSQVAPAE | 17.99 | YKGSQVAPA | 3895.2 |
| CB388-398 | CIKGPMVSKGY | 10.79 | IKGPMVSKG | 10309.3 |
| CB512-522 | RFVDSIPRNVT | 9.95 | FVDSIPRNV | 2842 |
| CB8-18 | VIYGPEPLHPL | 9.92 | IYGPEPLHP | 3346.6 |
| CB54-64 | EFFEATVLLAQ | 8.89 | FFEATVLLA | 2295.1 |
| CB95-105 | WYIGMIVAPVN | 8.54 | YIGMIVAPV | 6251.1 |
| CB495-505 | DYLAERVSHTK | 8.32 | YLAERVSHT | 9916.6 |
| CB480-490 | VVKQPGTEITA | 7.1 | VKQPGTEIT | 7368.1 |

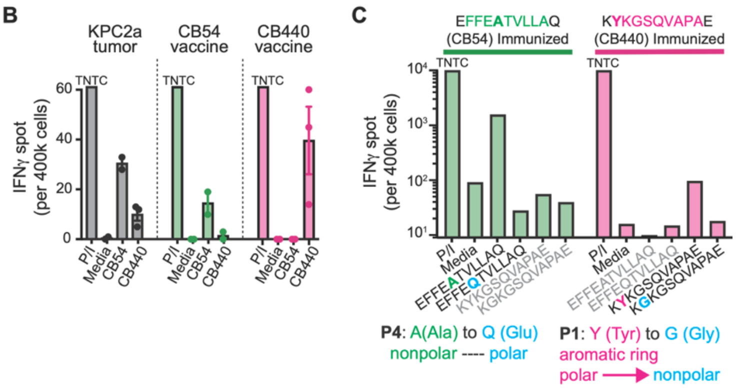


**Supplementary Figure 1. Identification and validation of I-A$^b$-restricted CBR peptides. Related to Figure 1. (A**) Candidate CBR peptides with predicted I-A$^b$ binding. Scores were determined using the custom Marc Jenkins algorithm and NetMHCII 2.1. **(B**) Number of IFN-γ-producing cells splenocytes cells isolated from day 7 *KPC*2a-bearing mice, CB54 TriVax-vaccinated mice (peptide + agonistic anti-CD40 + poly[I:C]), or CB440 TriVax-vaccinated mice. TNTC, too numerous to count indicates saturating IFNγ spots exceeding reliable quantification. **(C**) Number of IFN-γ-producing splenocytes cells from mice vaccinated with the indicated peptide and restimulated with same peptide, control peptides, or peptides that disrupt anchor residues position 4 (CB54) or position 1 (CB440).

#### Supplementary Figure 2

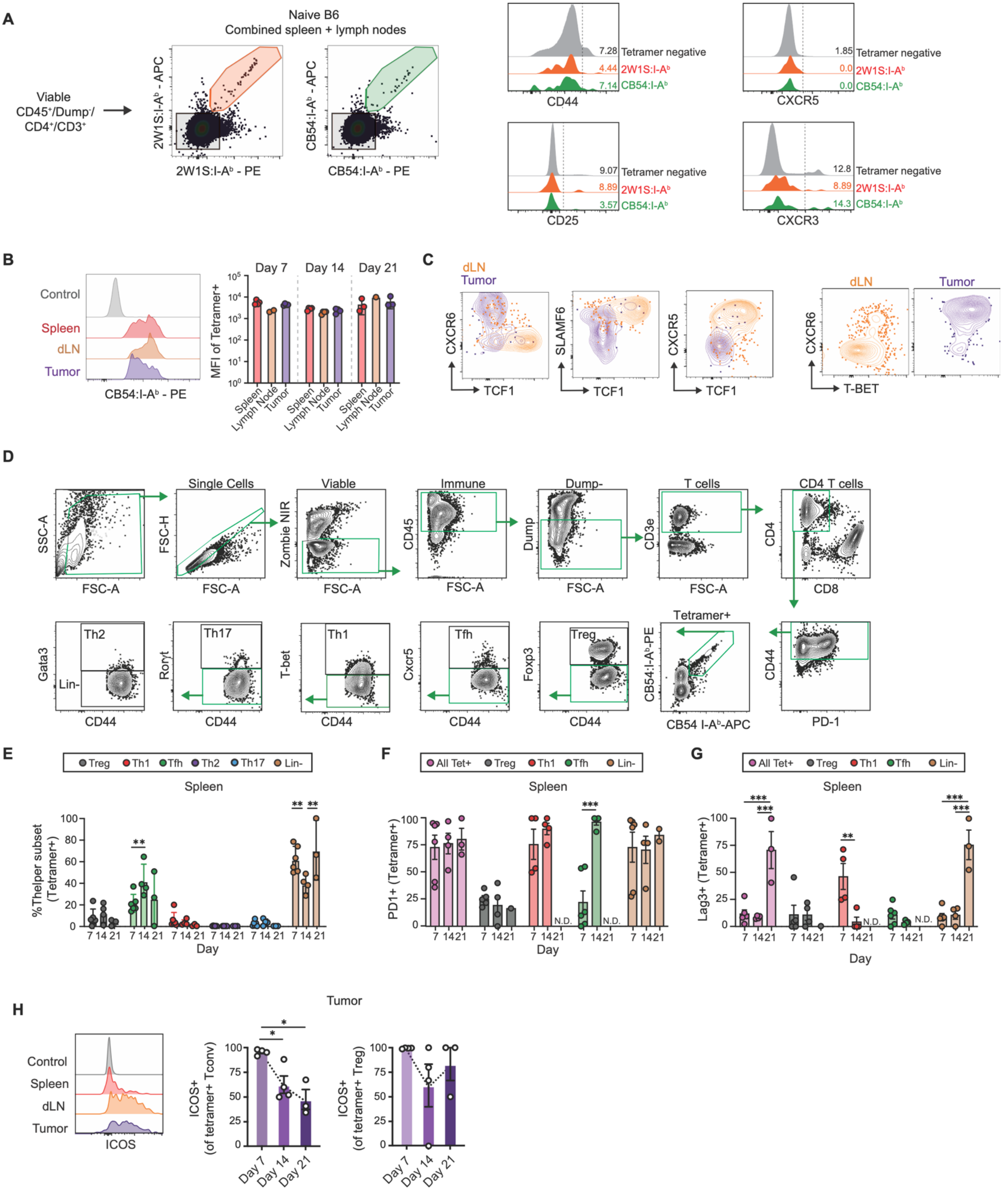

**Supplementary Figure 2. Fate of tumor-specific CD4 T cells during pancreatic tumor growth. Related to Figure 2. (A)** Representative flow cytometry plots and histograms gated on CB54:I-A$^b$-

specific and 2W1S:I-$A^b$-specific CD4 T cells in the pre-immune B6 repertoire. Plots are pooled from all secondary lymphoid organs from n=3 mice per group, following magnetic tetramer-based enrichment. (**B**) Representative histograms of CB54:I-$A^b$-tetramer staining in tetramer+ cells across tissues at day 14 post-tumor implantation (left) and tetramer MFI quantification at the indicated timepoints post orthotopic tumor implantation. Flow cytometry staining control (grey histogram), total splenic CD4 T cells. (**C**) Representative plots gated on CB54:I-$A^b$-specific CD4 T cells from the indicated tissues in orthotopic *KPC*2a-bearing mice. (**D**) Sequential gating strategy to delineate CB54:I-$A^b$-specific CD4 T cell differentiation state. $Lin^{-}$, lineage negative. (**E**) Frequency of CB54:I-$A^b$-specific CD4 T cells in spleen that adopt each T-helper differentiation state. (**F** and **G**) Frequency of splenic $tetramer^{+}$ CD4 T cell subsets at days 7, 14, and 21 that are PD-$1^{+}$ (F) and LAG-$3^{+}$ (G). Subsets comprising < 5 cells were excluded and are denoted N.D (not determined). **(H)** Representative histogram of ICOS staining across tissue on tetramer+ cells. **(I)** Frequency of Icos+ within tetramer+ Tconv (Foxp3-, left) or Treg (Foxp3+, right) CD4 T cells over time post-tumor. Data are pooled from n=3 naïve B6 mice for (A), each dot a mouse for (B, I), or reflect n=3 mice per group from 2 independent experiments (E-G). Data are represented as mean ± SEM. Statistical testing is one-way ANOVA with Tukey's posttest (B, E-G, I). *$p < 0.05$; **$p < 0.01$, ***$p < 0.001$.

#### Supplementary Figure 3

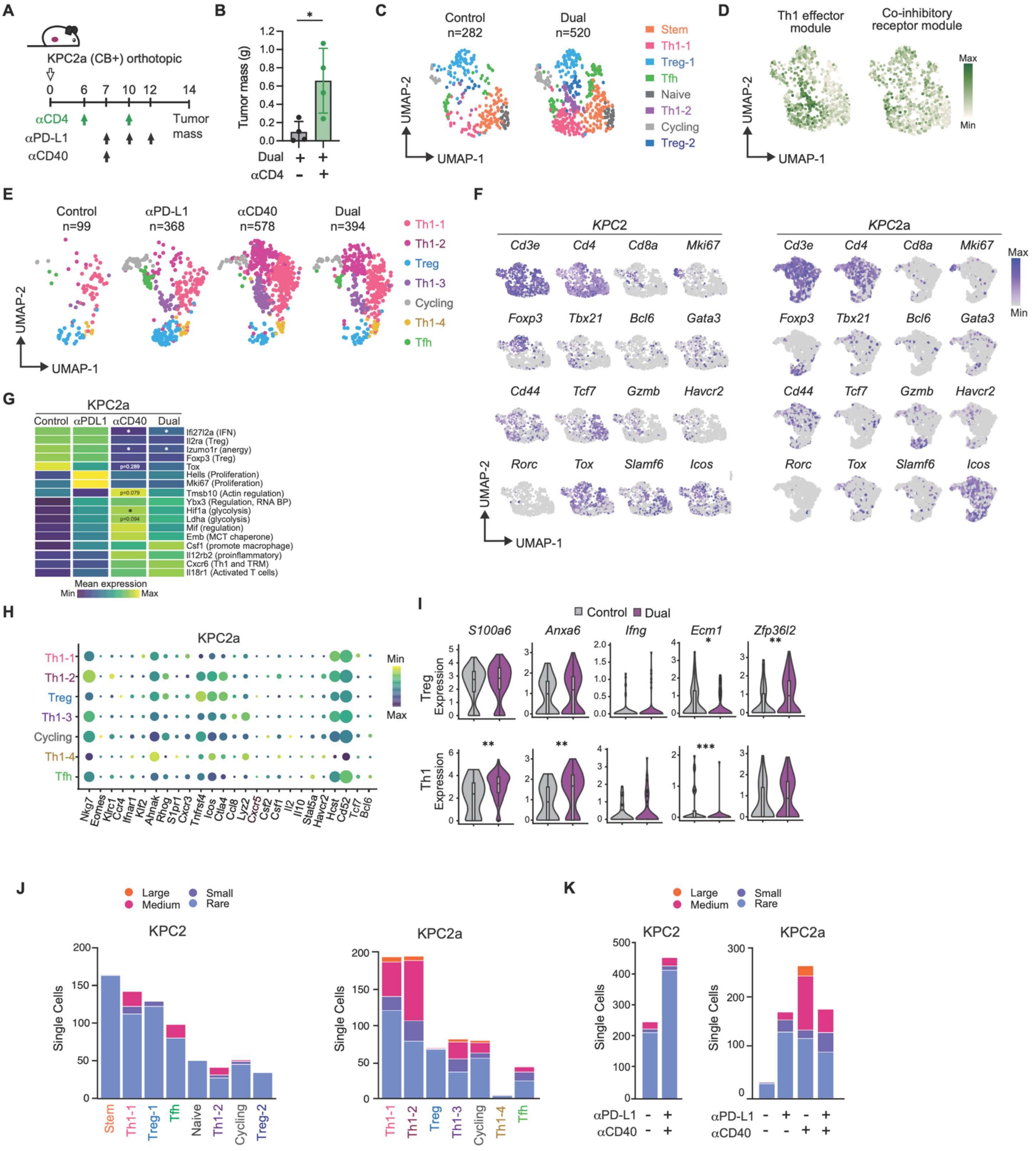

**Supplementary Figure 3. Single cell analyses of intratumoral CD4 T cells following agonistic αCD40 and αPD-L1. Related to Figure 3. (A**) Schematic to test the role of endogenous CD4 T cells at the effector phase of immunotherapy in the orthotopic *KPC*2a model. **(B**) Tumor weight in grams on day 14 from mice in (A). **(C**) UMAP split by treatment cohort of CD4 T cells in orthotopic *KPC*2 tumors and colored by cluster identity (resolution = 0.8). **(D**) Th1 effector module and co-inhibitory receptor module were visualized on merged UMAPs from (C). **(E**) UMAP plots split by treatment cohort of CD4 T cells in orthotopic *KPC*2a (CB+) tumors. Colored by cluster identity (resolution = 0.6). **F**) Feature plots of selected cluster defining genes on UMAP space from (C) and (E). **G**) Heatmap of selected differentially expressed genes on CD4 T cells from *KPC*2a tumors. Full DEG results for cross-therapy comparisons in Supplemental Table 1. **(H**) Dotplot of selected cluster defining DEGs in CD4 T cells from immunotherapy treated vs. control *KPC*2a dataset. **(I**) Violin plots of selected DEGs among Treg or Th1 cells in *KPC*2a tumors. **(J and K**) CD4 T cell clonal expansion was determined by scV(D)J-seq. Identical clonotypes were enumerated and classified as follows: large ($19<x\leq100$), medium ($4<x\leq19$), small ($2<x\leq4$), and rare ($0<x\leq2$). Number of single cells are visualized per cluster (J) or across treatment groups (K). All data are derived from n=3 mice pooled per treatment cohort per model. Data are presented as mean ± SEM, each dot a mouse. Unpaired Student's t test (B). Wilcoxon rank-sum tests with BH adjustment (G, I). *$p < 0.05$; **$p < 0.01$, ***$p < 0.001$

#### Supplementary Figure 4

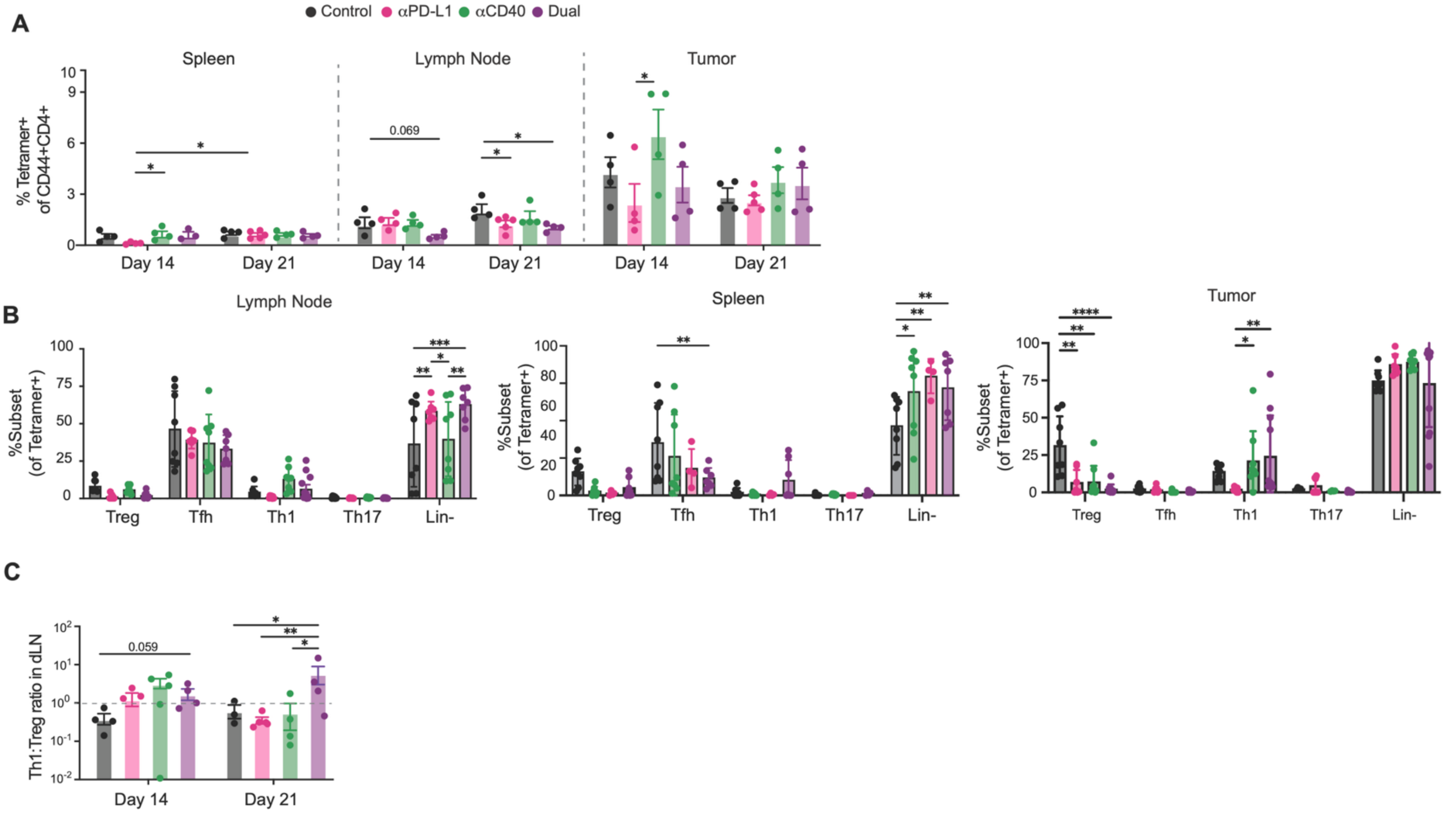


**Supplementary Figure 4. Anti-PDL1 promotes Tfh expansion whereas αCD40 promotes Th1 and transiently disrupts Treg cells. Related to Figure 4. (A)** Frequency of CB54-specific CD4+ T cells as a fraction of total antigen-experience cells (CD44+ CD4+) in the spleen, draining lymph node, and tumor across treatment groups, split by harvest at day 14 or day 21. **(B)** Frequency of CB54-specific CD4+ T cells differentiated to each of the T-helper lineages across therapies and tissues as gated in Figure S2D. **(C)** Intratumoral Th1 to Treg ratio quantified using numbers calculated in Figure 4I. Data are presented as mean ± SEM, with each dot representing one mouse reflect or n=3 mice per group from 2 independent experiments (B). Statistical analyses were performed using one-way ANOVA with Tukey's multiple comparison correction. *$p < 0.05$; **$p < 0.01$; ***$p < 0.001$; ****$p < 0.0001$.

### Supplementary Figure 5

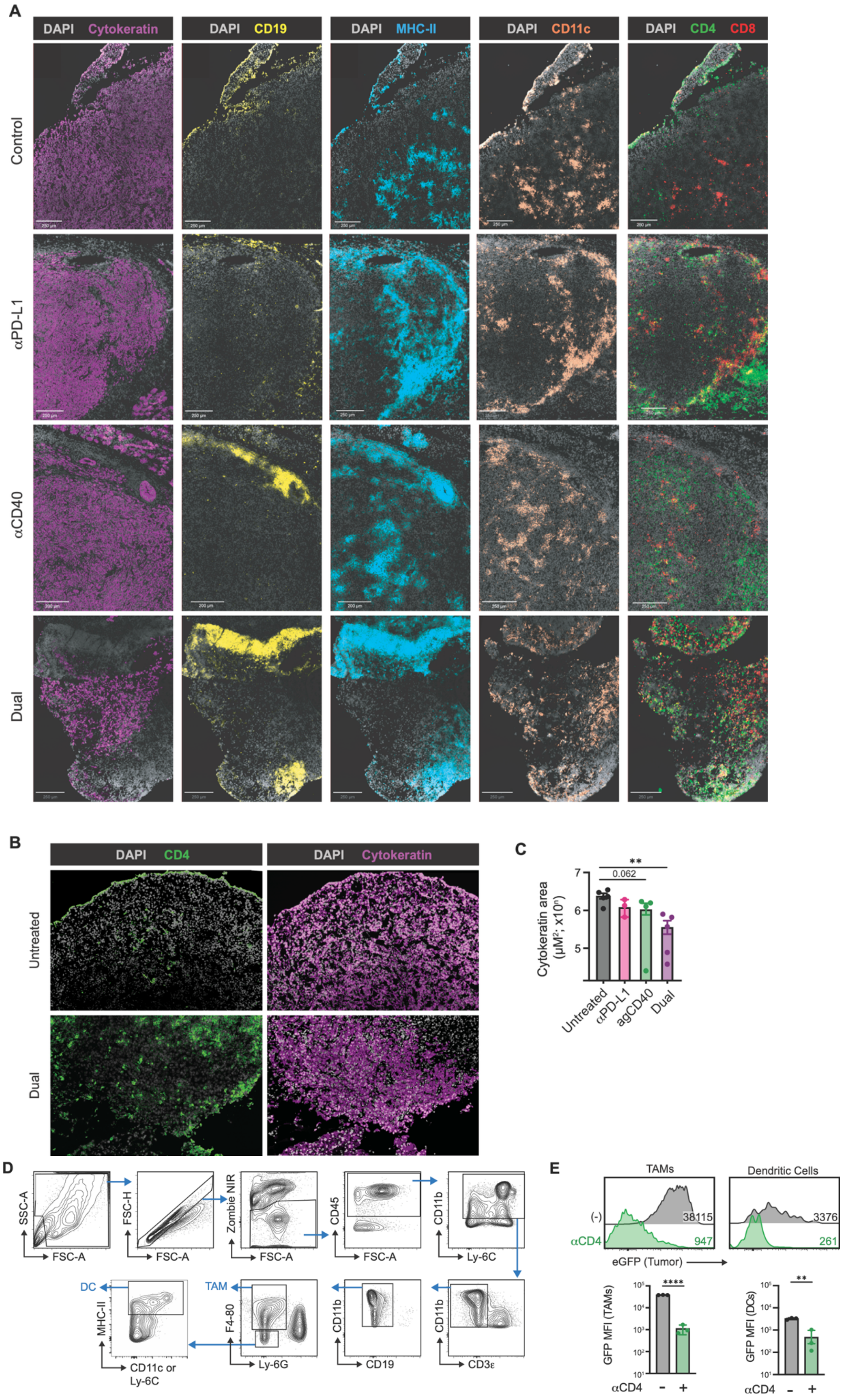

**Supplementary Figure 5. Multiplex immunofluorescence (IF) imaging of immune infiltration. Related to Figure 5.** (**A**) Representative multiplex immunofluorescence images of tumor sections stained for DAPI, cytokeratin, CD19, MHC-II, CD11c, CD4, and CD8. (**B**) Representative CD4, DAPI, and cytokeratin staining illustrating CD4 T cell infiltration into the tumor core. (**C**) Quantification of tumor (cytokeratin) area within total tissue ROI as measured by a Cytokeratin+ pixel thresholder in QuPath. **(D**) Gating strategy for intratumoral myeloid populations. **(E)** Representative histogram (top) of tumor antigen (GFP) loading in intratumoral TAMs and DCs as gated in (D) and quantification in isotype control or aCD4 depleted mice (bottom). Data are presented as mean ± SEM, each dot a mouse. one-way ANOVA with Tukey's multiple comparison correction for (C) and unpaired student t-test in (E). *$p < 0.05$; **$p < 0.01$, ***$p < 0.001$, ****$p < 0.001$

#### Supplementary Figure 6

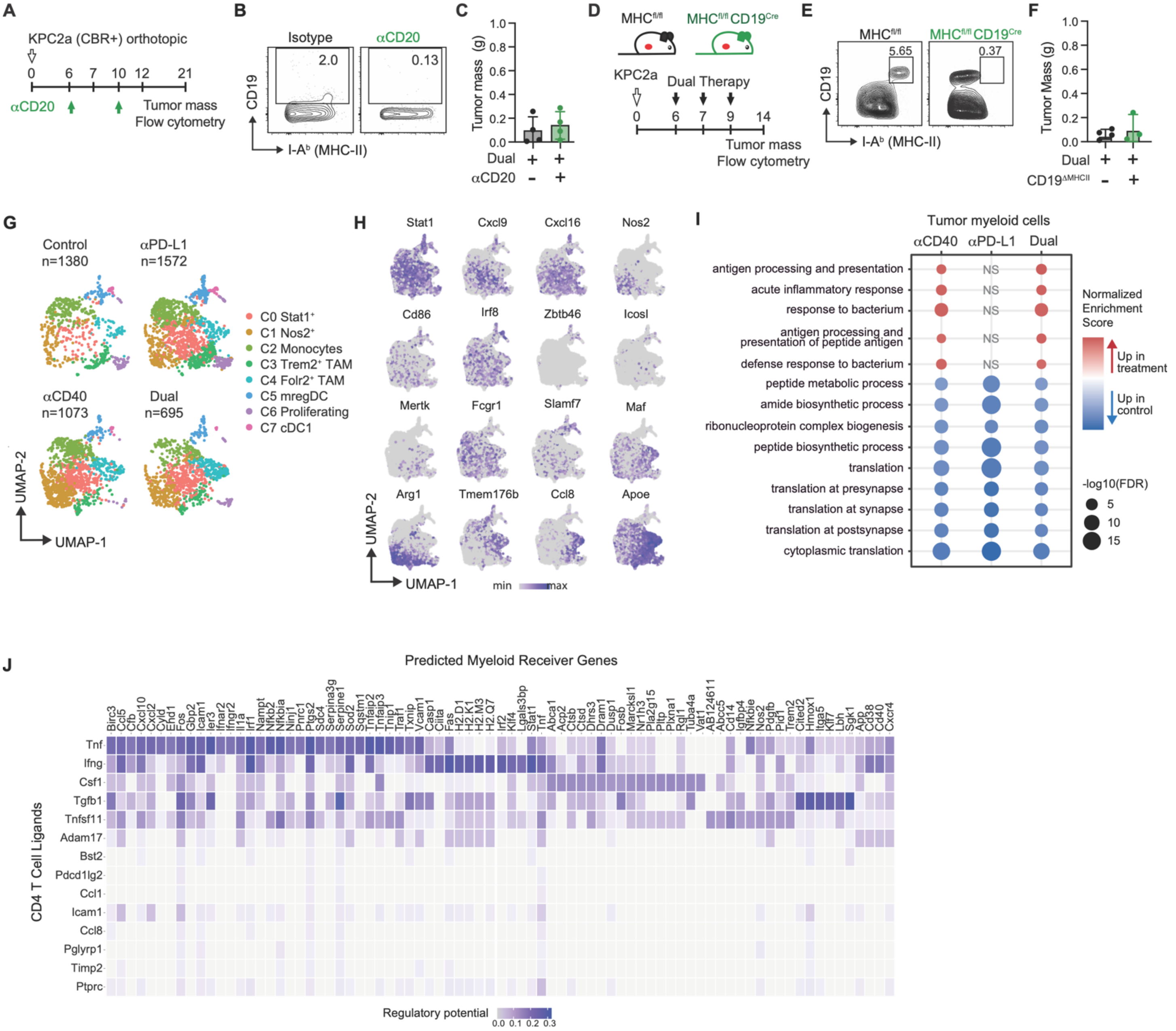


**Supplementary Figure 6, related to Figure 6. Immunotherapy alters myeloid cell fate and has antitumor effects independent of B cells. Related to Figure 6.** (**A**) Schematic of B cell depletion prior and during Dual therapy in orthotopic *KPC2a* tumor model. (**B**) Representative flow cytometry plot showing CD19 and MHC class II staining on on viable CD45$^{+}$ tumor infiltrating cells at day 21 post tumor implantation, showing depletion of B cells in anti-CD20-treated versus isotype control tumors at day 21. (**C**) Tumor weights from control and anti-CD20-treated mice. (**D**) Schematic of orthotopic *KPC*2a tumor implantation in control (MHC-II$^{fl/fl}$) or mice that lack MHC-II on B cells (MHC-II$^{fl/fl}$ x CD19$^{Cre+}$) mice. (**E**) Representative plots gated on viable CD45$^{+}$ tumor-infiltrating cells showing MHC-II depletion on B cells of MHC-II$^{fl/fl}$ CD19$^{Cre+}$ mice on day 14 post-implantation. (**F**) Tumor weights from mice in (E) on day 14. (**G**) UMAP of myeloid cluster identity split by treatment group, with cell numbers per group indicated, colored by cluster identity. (**H**) Feature plots of select cluster-defining myeloid marker genes. (**J**) NicheNet heatmap predicting CD4 T cell-derived ligands (rows) that drive the Dual-therapy associated DEG transcriptional response of myeloid cells (columns).

Myeloid receiver genes inputted to NicheNet were identified by comparing untreated and Dual therapy myeloid cells via Wilcoxon rank-sum tests with BH adjustment (Seurat FindMarker). Expressed genes on CD4 T cells (senders) were used to calculate regulatory potential, an estimate of how likely a ligand is to regulate the expression of a target gene through known signaling and gene regulatory networks. Data are presented as mean ± SEM, each dot a mouse in (C) and (F). Unpaired Student's t-test for (C) and (F).

#### Supplementary Figure 7

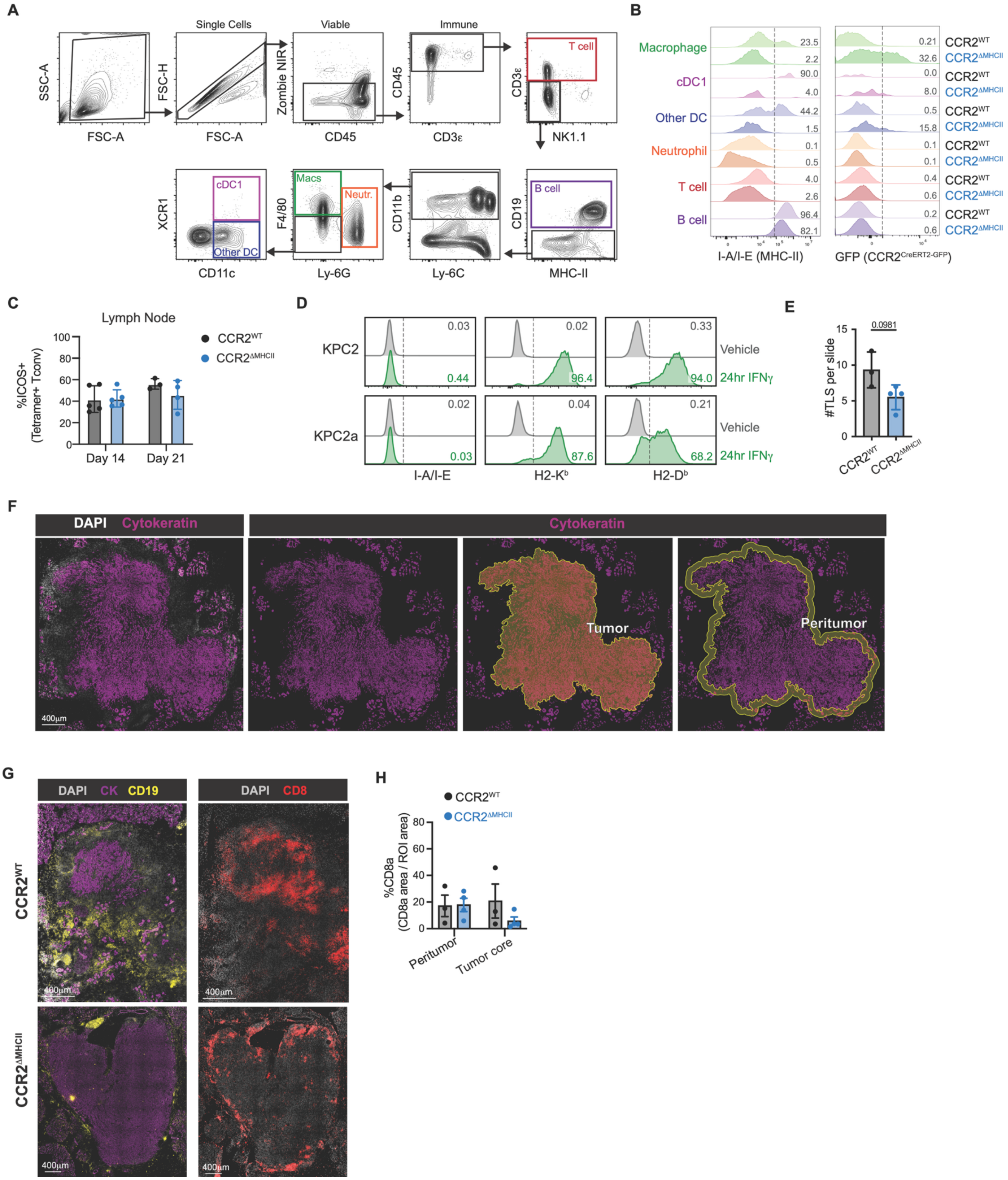


**Supplementary Figure 7, related to Figure 6. Immunotherapy requires antigen presentation by myeloid cells CD4 T cell infiltration, TLS formation, and antitumor effects Related to Figure 6.**

(**A**) Gating strategy for identification of immune cell subsets from splenocytes from a $CCR2^{WT}$ mouse. This strategy permits gating on DCs independent of MHC-II. (**B**) Histograms of MHC-II and GFP (Cre recombinase) gated on the indicated immune subsets in spleen from tamoxifen-treated mice of the $CCR2^{WT}$ or $CCR2^{\Delta MHCII}$ genotype. (**C**) Frequency of dLN-residing tetramer$^{+}$ CD4 $T_{conv}$ cells that express ICOS in tamoxifen-treated $CCR2^{WT}$ versus $CCR2^{\Delta MHCII}$ mice. (**D**) Histograms of I-$A^{b}$, H2-$K^{b}$, and H2-$D^{b}$ on *KPC* tumor cells treated with or without 24 h treatment with vehicle (media) or recombinant mouse IFN-γ *in vitro*. (**E**) Number of TLS per slide. (**F**) Representative IF staining demonstrating tumor periphery versus tumor core. (**G**) Localization of CD8 T cells in day 14 orthotopic *KPC*2a tumors from tamoxifen-treated $CCR2^{WT}$ and $CCR2^{\Delta MHCII}$ mice. (**H**) Percent area containing CD8 T cells in tumor periphery and tumor core. Data are mean ± SEM and each dot is an independent mouse or tumor (C, E, H). Unpaired student's t test (E).

#### Supplementary Figure 8

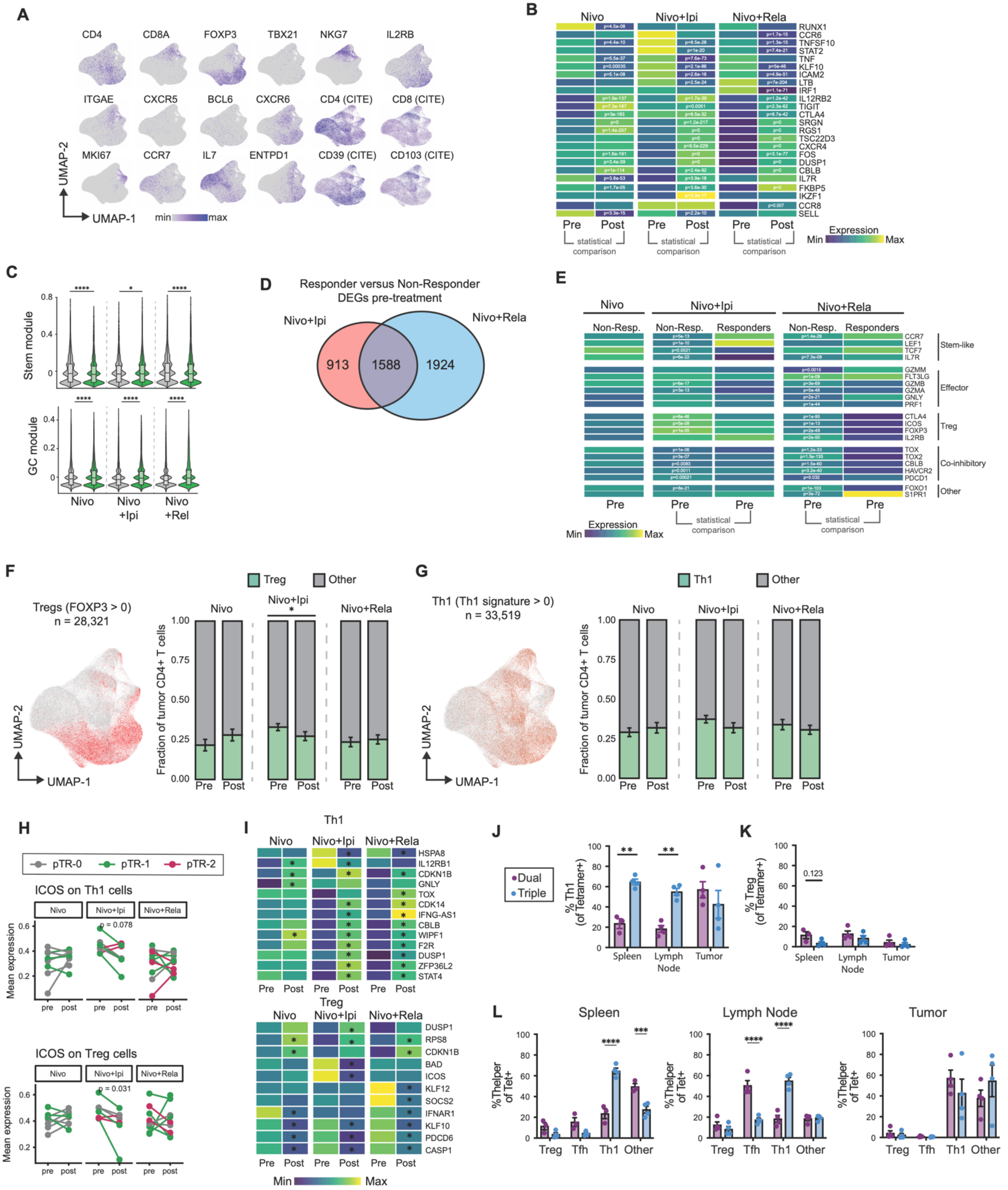

**Supplementary Figure 8. Extended analysis of CD4+ T cell transcriptional responses to immunotherapy in human HNSCC and IL-15C therapy effect in murine CD4 T cells. Related to Figure 7.**

**(A)** Feature plots of cluster-defining genes and CITE-seq markers projected onto UMAP space as in Figure 7A. **(B)** Extended heatmap of DEGs between pre- and post-treatment, split by treatment arm. **(C)** Violin plots of stem and germinal center (GC) gene module scores before and after immunotherapy across treatment cohorts. **(D)** Venn diagram of DEGs in responders versus non-responders for nivolumab + ipilimumab (red) and nivolumab + relatlimab (blue), with shared DEGs shown in the overlap (gray). **(E)** Heatmap of select DEGs between responders (pTR-2) and non-responders (pTR-0 and pTR-1). **(F)** Feature plot of cells identified as Treg (left) and stacked bar plot quantification of Treg frequency pre- and post-treatment, split by treatment arm (right). **(G)** Feature plot of cells identified as Th1 (defined by the Th1 signature described in STAR Methods) (left) and stacked bar plot quantification of Th1 frequency pre- and post-treatment, split by treatment arm (right). **(H)** Patient-level paired pre- and post-treatment mean ICOS expression in Th1 (top) and Treg (bottom) cells, split by treatment arm. Connected dots represent individual patients, colored by pTR histopathology. **(I)** Heatmap of pre- versus post-treatment DEGs in Th1 (top) and Treg (bottom) cells. **(J and K)** Frequency of Th1 (J) and Treg (K) among tetramer+ CD4+ T cells across tissues in Dual versus Triple therapy at day 14. **(L)** T helper subset frequency among tetramer+ CD4+ T cells across tissues in Dual versus Triple therapy at day 14, shown for spleen (left), draining lymph node (middle), and tumor (right).

Full DEG results for responder versus non-responder analysis (E) and DEGs across therapy for Th1/Treg (I) in Supplemental Table 4. Statistical analyses were performed using two-sided Wilcoxon rank-sum tests with BH correction for (C), Seurat FindMarkers for (D), (E), and (I), chi-square tests with adjustment for (F) and (G), paired t tests for (H), and one-way ANOVA with Tukey's multiple comparison correction for (J–L). $^{*}p < 0.05$; $^{**}p < 0.01$; $^{***}p < 0.001$; $^{****}p < 0.0001$.
